# Cell cycle-dependent protein dynamics in budding yeast resolved by deconvolution of bulk proteomics

**DOI:** 10.64898/2026.02.12.705502

**Authors:** Andre Zylstra, Mattia Rovetta, Silke R. Vedelaar, Christian Bleischwitz, Julius A. Fülleborn, Yulan van Oppen, Hugo P. Markus, Kerlen T. Korbeld, Andreas Milias-Argeitis, Katarzyna Buczak, Alexander Schmidt, Matthias Heinemann

## Abstract

The cell division cycle is characterised by oscillatory dynamics in regulatory mechanisms and biosynthesis, coordinated with genome replication and segregation. To understand these dynamics, quantitative cell cycle-dependent protein concentration data is essential. Unfortunately, accurate resolution of cell cycle-dependent protein dynamics is challenging because single-cell proteomics is currently infeasible and bulk proteomics requires inherently imperfect cell synchronisation. Here, we developed a computational method to deconvolve cell cycle-dependent protein concentration dynamics and applied it to new budding yeast bulk proteome data. Key to this method was a yeast population model, parameterised with experimental cell cycle progression and volume growth data, for quantifying the desynchronisation in sampled populations. We performed deconvolution on 3373 proteins, using cross-validation to determine regularisation parameters, and identified 563 proteins with cell cycle-dependent dynamics. Many of these dynamics were consistent with known yeast biology and dynamic proteins were enriched for several metabolic process, extending previous observations and supporting the emerging picture of metabolic activity as varying substantially over cell cycle phases. We consider the generated cell cycle-resolved budding yeast proteome data a key resource.

## Introduction

Successful completion of a eukaryotic cell cycle requires cooperation between diverse cellular processes. While the dynamics of the cell cycle control and DNA replication machinery have been long studied, it was recently found in budding yeast that lipid, carbohydrate, and protein synthesis are partially temporally segregated across the cell cycle, giving rise to coordinated metabolic oscillations (Takhaveev *et al*, 2023). Because such biosynthetic dynamics contribute to cell cycle control (Litsios *et al*, 2019), it is important to uncover the mechanisms that give raise to this temporal segregation. To answer this question, and many other cell cycle related questions, it will be essential to have a detailed map of protein dynamics over the cell cycle.

Unfortunately, our current knowledge of cell cycle-dependent protein concentrations is still very limited. Ideally, cell cycle-dependent protein concentrations would be measured dynamically in single cells, for instance with time-lapse microscopy. Recently, Litsios and colleagues achieved a major step towards this goal by using high-throughput microscopic analyses with GFP-tagged proteins to assay >2800 proteins during six cell cycle phases (Litsios *et al*, 2024). However, this analysis was based on snapshots rather than time lapses, and as such provided limited and uneven cell cycle resolution; for instance, three data points were collected during the short M phase (metaphase, anaphase, and telophase) while there was only one for S/G2, which is longer. In addition, fluorescence microscopy can suffer from artifacts stemming from incomplete fluorophore degradation (Khmelinskii *et al*, 2016) and potential disturbances of the natural protein degradation dynamics by the fluorophore tag (Koren *et al*, 2018; Lin *et al*, 2018). Furthermore, microscopic analyses would also not allow for determination of post-translational protein modifications, whose mapping will likely also be needed for the full picture of cell cycle-dependent protein dynamics.

Thus, at least in the near future, population-level proteomics techniques applied to synchronised cell populations will still be necessary to achieve this aim. Proteomic analyses of synchronised populations have previously been performed with budding yeast using alpha factor arrest-release (Kelliher *et al*, 2018; Campbell *et al*, 2020; Flory *et al*, 2006), inducible cyclin expression (Zhang *et al*, 2019), or centrifugal elutriation (Blank *et al*, 2020). However, due to incomplete arrest or heterogenous release, synchronisation is never perfect and, even if it were, intrinsic cellular stochasticity would still lead to progressive desynchronisation over time. In budding yeast, asymmetric cell division and the longer G1 phases of newborn cells (Hartwell & Unger, 1977) make the issue of desynchronisation even worse. Unfortunately, desynchronisation blurs concentration dynamics measured at the population level, leading to an underestimation of true cell cycle–dependent protein dynamics. Thus, even with experimental synchronisation techniques, proteomics provides only a limited view of true protein dynamics.

Given that experimental synchronisation is inherently imperfect, a different approach is required to determine the true cell cycle-resolved protein dynamics in budding yeast. Computational deconvolution of data obtained from partially synchronised cultures is a potential solution. Such deconvolution solves the ‘inverse problem’ of reconstructing the underlying cell cycle-dependent protein concentration dynamics from the observed bulk signal, given information about the distribution of cells across cell cycle phases in the sampled culture. To the best of our knowledge, in budding yeast, deconvolution has so far only been applied to transcriptome data (Rowicka *et al*, 2007; Orlando *et al*, 2007; Guo *et al*, 2013; Bar-Joseph *et al*, 2004; Qiu *et al*, 2006).

In this work, we developed and applied a computational framework that estimates cell cycle-dependent protein concentrations with high cell cycle resolution from population-level proteomics data obtained from partially synchronised populations. Our approach involved estimation of the convolution kernel from simulated cell populations using a parameterised cell cycle model. We modelled the bulk protein concentration measurements as mixtures of cell cycle stage-specific concentrations and performed deconvolution using a regularised least squares method incorporating cell cycle periodicity constraints. Using samples from elutriated cultures, we measured protein concentrations by mass spectrometry proteomics, applied our deconvolution approach to 3373 quantified proteins, and identified 563 proteins with higher confidence cell cycle-dependent dynamics. We found that the deconvolved dynamics of these proteins were consistent with known yeast cell cycle biology, with respect to cell cycle-dependent metabolic dynamics and imputed transcription factor regulatory activity, supporting the validity of the deconvolved cell cycle-dependent protein concentration trajectories.

As such, here we make two contributions: (i) an approach that can be applied to estimate cell cycle-dependent concentration dynamics from bulk proteomics data and (ii) the cell cycle-resolved protein concentration dynamics of budding yeast as a resource. We envision that this dataset will be used to answer urgent cell cycle related questions, such as how the recently observed temporal segregation of biosynthetic processes emerges and how this exerts cell cycle control.

## Results

To develop our computational deconvolution approach, wherein we would estimate an underlying cell cycle-dependent protein concentration dynamic from population-level experimental measurements, we had to solve three main challenges. First, we had to derive an equation which represented the convolution process, relating an underlying protein concentration dynamic we wished to infer to the population-level measurements obtained in an experiment. Importantly, this convolution equation would necessarily include a convolution kernel, representing the ‘blurring’ effect of cell cycle desynchronisation in the population. Second, we had to develop a model of yeast population cell cycle progression and volume growth which enabled us to estimate the specific convolution kernel corresponding to each experiment whose population-level measurements we wished to deconvolve. Lastly, we required an algorithm, which could perform the deconvolution given population-level measurement data and the estimated convolution kernel as inputs, i.e. solving the convolution equation. Critically, this deconvolution algorithm would have to be robust to realistic amounts of measurement noise and imprecision in the convolution kernel estimate.

### Derivation of the convolution equation

To allow expressing protein concentrations as a function of cell cycle progression, it was useful to express these dynamics in terms of an abstract cell cycle progression axis rather than in terms of time since individual cell cycles have variable durations. We defined a unitless cell cycle progression axis, *φ*, over the domain [-1, 1], where *φ* = −1 corresponded to cell birth (i.e. the moment of cytokinesis where bud and mother become separate cytoplasmic compartments), *φ* = 0 to START of a cell cycle, and *φ* = 1 to START of the next cell cycle [Fig. 1A]. Mother and daughter early-G1 phases correspond to separate parts of the *φ* axis, but the rest of the mother and daughter cycles map to a shared region of *φ*. A full mother cycle, running from START to START, occurs over 0 ≤ *φ* < 1. A complete daughter cycle runs from birth/cytokinesis at *φ* = −1 to the subsequent cytokinesis at *φ*_*cyt*_, where 0 < *φ*_*cyt*_ < 1 and the specific *φ*_*cyt*_ value was calculated based on experimental cell cycle phase duration data.

**Figure 1.**
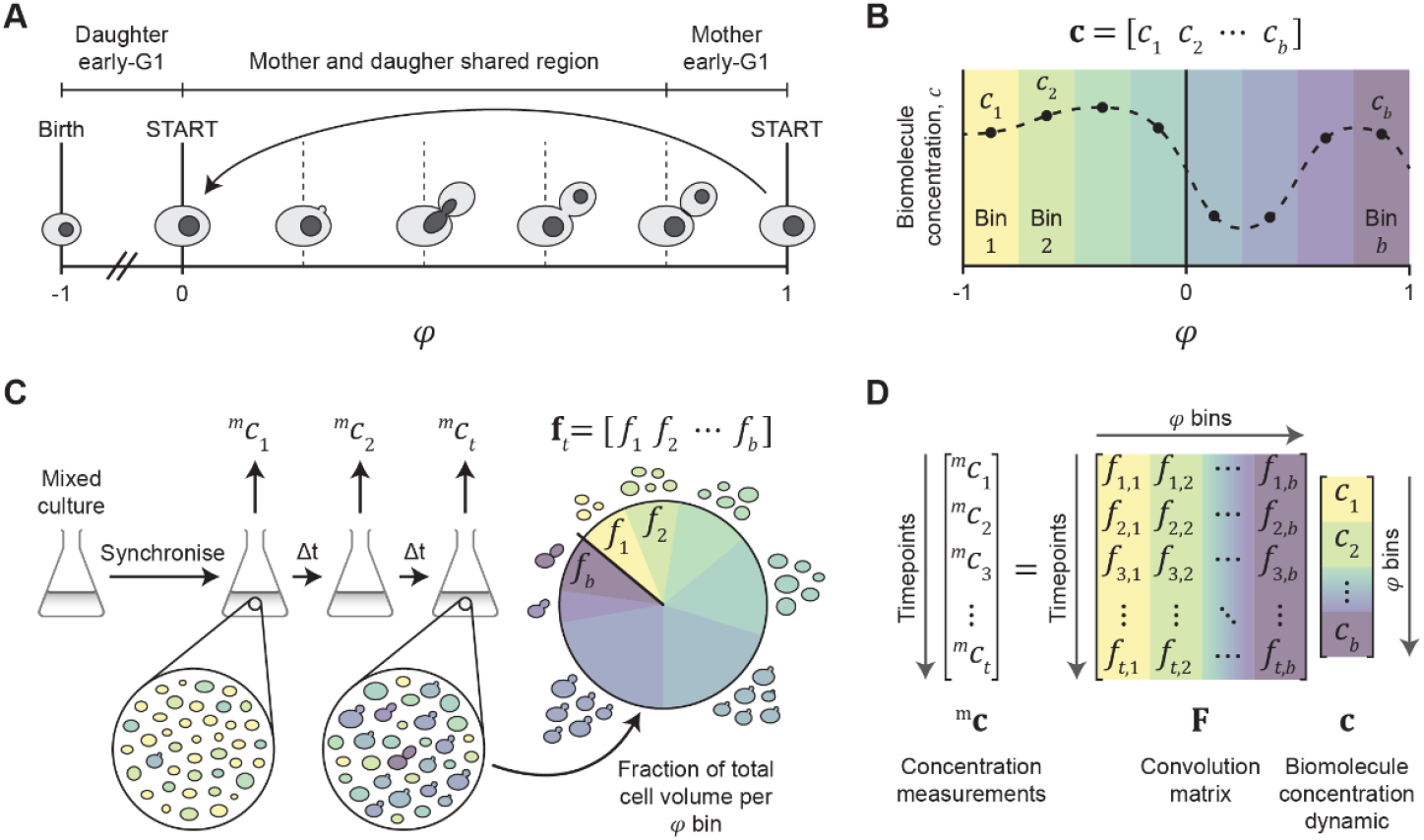
A linear convolution model describes the influence of cell cycle desynchronisation on population-level measurements. **A**. *φ* represents a unitless cell cycle progression axis over the domain [-1, 1] onto which cell cycles of different durations and stages of progression are mapped. *φ* = −1 corresponds to cell birth (i.e. the moment of cytokinesis where bud and mother become separate cytoplasmic compartments), *φ* = 0 to START of a cell cycle, and *φ* = 1 to START of the next cell cycle. A daughter cell’s position begins at *φ* = −1 and increases up to *φ*_*cyt*_, the value of *φ* at cytokinesis, where 0 < *φ*_*cyt*_ < 1. For mother cells, upon reaching START at *φ* = 1, the *φ* position immediately shifts to *φ* = 0 such that mother cells are always between 0 and 1. **B**. The mean (over many single cells) cell cycle-dependent concentration of a protein, *c*, is a function over *φ*. By discretising the *φ* axis into *b* equally spaced bins, the cell cycle-dependent concentration is represented as a vector **c** with *b* elements. **C**. Measurements of a protein’s concentration are collected in a time course experiment, ^*m*^*c*_1_, ^*m*^*c*_2_ ⋯, ^*m*^*c*_*t*_, comprising the vector ^*m*^**c**. The cell cycle desynchronisation in the sampled population at time point *t* is reflected in the vector **f**_*t*_, i.e. the fractional distribution of cell volumes across the *φ* bins at time point *t*. **D**. The relationship between **c** and ^*m*^**c** for a whole time course experiment is expressed as the convolution equation ^*m*^**c** = **Fc**. The convolution matrix **F**, with row vectors **f**_1_, **f**_2_ ⋯, **f**_*t*_, represents the effect of population-level desynchronisation on the set of concentration measurements.

We discretised the *φ* axis into *b* equally spaced bins, meaning that we could describe cell cycle-dependent protein concentration dynamics as vectors with *b* elements. Due to the intrinsic stochasticity of biochemical processes, individual cells at the same *φ* position can differ somewhat with respect to protein concentrations. Nonetheless, by taking the mean over many cells at each *φ* position, we would obtain the concentration dynamic representative of a single cell’s behaviour. As such, we could represent the mean cell cycle-dependent single cell concentration dynamic of a protein as a column vector **c** of length *b* [Fig. 1B].

Ideally, we would like to measure **c** directly from population-level measurements, but we cannot achieve that due to imperfect synchronisation methods and inherent progressive desynchronisation of the cell population over time. Instead, to estimate **c** from population-level measurements by deconvolution, we required a convolution equation relating **c** to the population-level experimental data via the convolution kernel, which represented the desynchronisation process. To derive this convolution equation, we considered a typical time course experiment, which would start with a partially synchronised population (e.g. synchronised by elutriation) from which we would take measurements. At the *t*-th time point, we would take a sample, measure the abundance of the protein of interest, and then divide this abundance by the total cell volume in the sample to obtain measured population-averaged concentration, ^*m*^*c*_*t*_. The binned concentration values in vector **c** would each contribute to ^*m*^*c*_*t*_ proportional to the fraction of the sampled cell volume that came from cells belonging to each *φ* bin [Fig. 1C]. We represented these volume fractions, at the *t*-th time point, as a row vector **f**_*t*_ of length *b*. Thus, a population-level measurement ^*m*^*c*_*t*_ at time point *t* would resemble the dot product of the two vectors **f**_*t*_ and **c**, ^*m*^*c*_*t*_ = **f**_*t*_ **· c**.

Each measurement time point in a time course experiment would give rise to a similar equation. If we collected a total of *n* measurements over a time course experiment, we could represent these measurements as a column vector ^*m*^**c** of length *n*. Cells would grow, progress in their cell cycles and potentially divide between time points. Hence, there would be *n* different **f**_*t*_ vectors, one for each measurement time point. We constructed the *n* × *b* matrix **F** by stacking these vectors row-wise and thereby obtained a single expression representing the whole experiment, ^*m*^**c** = **Fc** [Fig. 1D]. The matrix **F** encodes a discretised version of the convolution kernel which ‘blurs’ the values of a cell cycle-dependent protein concentration dynamic into the population-level concentration values measured in an experiment. The ‘blurring’ process is a so-called convolution operation, so we refer to **F** as a convolution matrix hereafter.

We now had a mathematical relationship between the cell cycle-dependent concentration dynamic for a protein of interest, **c**, and population-level measurements taken during a time course experiment, ^*m*^**c**. To be able to solve for the unknown vector **c**, i.e. to deconvolve the measurement data, we required an accurate estimate of the convolution matrix **F**, which would be unique for each time course experiment.

### Modelling yeast population growth

To estimate the convolution matrix **F** for a given experiment, we needed to know, for each experimental time point, what proportion of the sampled cell volume came from cells belonging to each *φ* bin. To obtain this information, we developed a computational model with which we could simulate the temporal evolution of an initially synchronised yeast cell population. Specifically, we simulated a population consisting of individual single cells, wherein each cell cycle of every cell was defined by a set of parameters: an initial volume (i.e. the volume at cytokinesis of the previous cell cycle), two linear volume growth rates (for pre- and post-budding), and cell cycle phase durations. Though cell separation is not a cell cycle phase *per se*, we also included the duration between cytokinesis and cell separation to enable comparison of simulations with experimental data where unseparated mothers and daughters were detected as single particles. To simulate heterogeneity between cell cycles of the modelled population, we randomly sampled parameters for each cell cycle from multivariate parameter probability distributions. We ensured that these distributions reflected realistic cycle-to-cycle variability in volume growth and cycle phase progression by fitting them to experimental data.

We obtained empirical distributions for the required cell cycle parameters from time-lapse imaging of cells grown on microfluidic devices in 2% glucose minimal medium. We estimated cell volumes from segmented cell areas, obtained with the ImageJ plugin BudJ (Ferrezuelo *et al*, 2012), assuming ellipsoid cell morphology. From these volumes, we found the birth volumes of newborn daughter cells, as well as volume growth rates in both the G1 and S/G2/M phases for mothers and daughters, assuming linear growth in these two phases [Fig. 2A]. To pinpoint key cell cycle events, we used a strain that had the histone H2A tagged with sfGFP (Hta2-sfGFP) and the cell cycle inhibitor Whi5 tagged with mCherry (Whi5-mCherry). With this strain, we identified START and mitotic exit, respectively, from the moments where Whi5-mCherry exited and re-entered the nucleus. Using the Hta2-sfGFP marker, we identified karyokinesis as the moment when nuclei started to separate before division. Lastly, we determined the moments of budding, cytokinesis and separation from the brightfield images. We then found the durations of the cell cycle phases between all these events [Fig. 2B].

**Figure 2.**
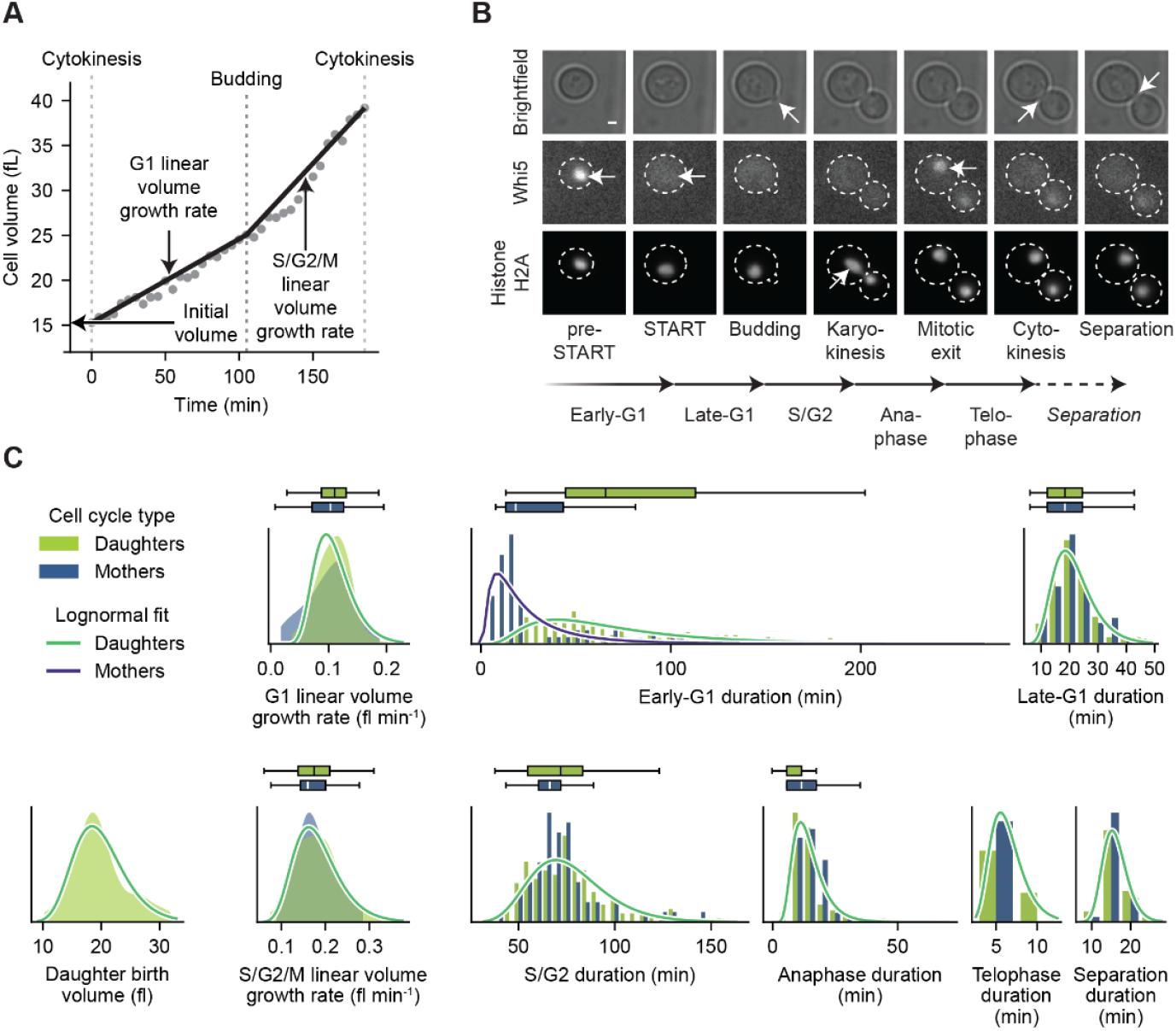
Parameters for cell cycle phase durations and volume growth. **A**. A model for cell cycle-dependent cell volume growth. The initial volume is the volume immediately following cytokinesis at the end of the preceding cell cycle. Grey dots indicate the measured volumes for an example cell cycle. The G1 linear volume growth rate is the gradient of the line between the initial volume and the volume at budding. The S/G2/M linear volume growth rate is the gradient of the line between the volume at budding and the volume at the subsequent cytokinesis. **B**. Key cell cycle events and the durations between them were identified from time lapse fluorescence microscopy of an Hta2-sfGFP Whi5-mCherry strain growing on a microfluidic device. White dotted lines indicate the cell and bud boundaries. White arrows highlight the defining feature of each cell cycle event. Scale bar is 1 µm. **C**. Distributions of cell cycle phase duration, initial volume, and volume growth parameters were determined from pooled data from 3 time lapse fluorescence microscopy experiments with cells grown on microfluidic devices. “Daughter” cell cycles were the first cell cycle undergone by a cell after separating from its mother (n=134). “Mother” cycles were any subsequent cell cycle (n=58).

Having determined cell cycle phase duration and volume growth dynamics, we calculated multivariate parameter distributions to allow sampling parameters either for simulated daughter or mother cell cycles. The distributions for volume growth rates and cell cycle phase durations were generally similar between mothers and daughters [Fig. 2C]. However, as reported previously (Hartwell & Unger, 1977), the early-G1 durations of daughter cells (median 60.0 min, IQR 65.0 min) were typically much longer compared with mothers (median 15.0 min, IQR 28.75 min) [Fig. 2C upper middle panel]. As the distributions were right-skewed, we estimated the distribution for each single parameter as a log-normal distribution by maximum likelihood estimation [Fig. 2C, Supp. Table 1]. Then, to obtain the multivariate log-normal distribution for daughter cycles, distribution D, we took the log-means of the single parameter distributions and calculated the covariance matrix from the log-standard deviations and the correlation matrix between log-transformed parameters [Supp. Fig. 1, Supp. Table 2]. Since the mother and daughter parameter distribution were similar, we also calculated multivariate distribution M (for mothers) using daughter cell data and correlations, except that we included the shorter mother early-G1 data instead.

Having determined the parameter distributions describing cell cycle phase durations and volume growth in single cells, we built the computational model simulating the temporal development of a yeast population in liquid culture, initially synchronised by elutriation. In the model, we accounted for cell volume growth, cell division, cell-to-cell heterogeneity in terms of cell growth and cell cycle progression, as well as the differences between daughter and mother cells. A simulated cell population comprised hundreds of thousands of cells undergoing their cell cycles. Each cell cycle was represented as a set of parameters, consisting of an initial volume (*v*_0_), two linear volume growth rates (*g*_1_ in G1 and *g*_2_ in S/G2/M), an initial time (*t*_0_, i.e. the time in the simulated experiment when the cycle began), and cell cycle phase durations (*d*_1_–*d*_6_) [Fig. 3A]. There were two distinct successive procedures involved with producing a simulated population: first, we generated the first set of cell cycles for newborn cells that represented the initial population of elutriated cells. Then, whenever a cell cycle ended, we generated a subsequent mother cell cycle and a new daughter cell, thereby continuing the population’s development until the end of the experiment.

**Figure 3.**
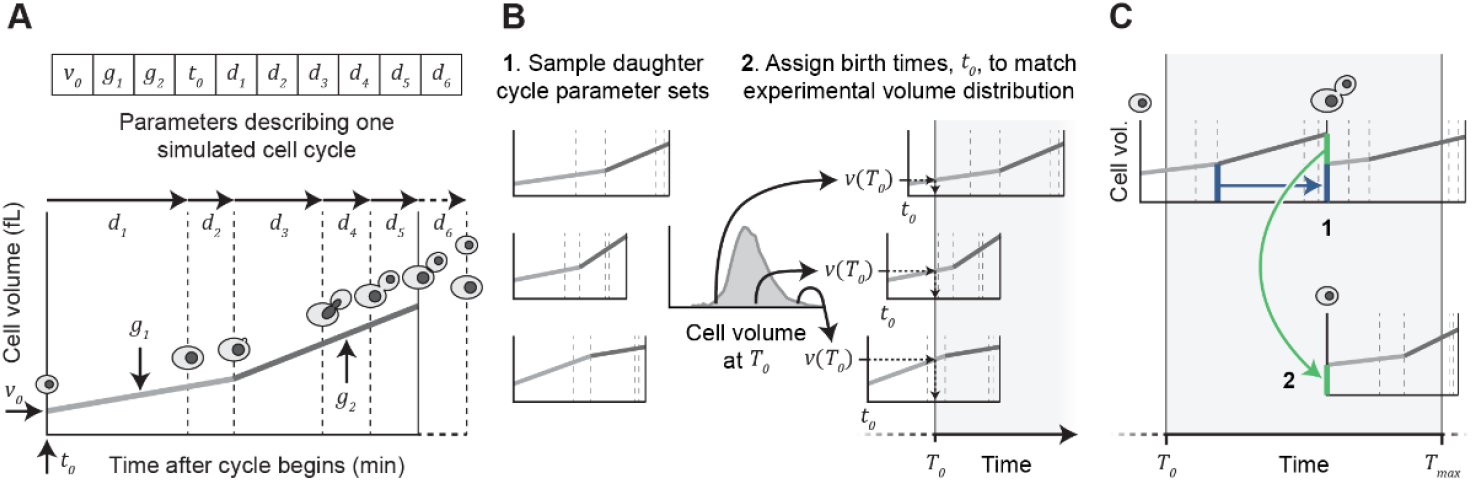
Parameterisation of single cell cycles within a simulated population. **A**. A single simulated cell cycle is represented by 10 parameters. The cycle initial volume is *v*_0_ and the cell grows with linear volume growth rate *g*_1_ during G1 and *g*_2_ during S/G2/M. *t*_0_ is the moment of birth/cytokinesis of the preceding cell cycle and the parameters *d*_1_, *d*_2_, *d*_3_, *d*_4_, and *d*_5_ indicate the durations of early-G1, late-G1, S/G2, anaphase, and telophase respectively. The separation duration, *d*_6_, is not part of the cell cycle, but is included in the model to enable comparison of simulated population volume distributions with those obtained from instruments which record unseparated mother-daughter pairs as single particles. Most parameters for each simulated cell cycle were drawn from multivariate log-normal distributions derived from the data measured from cells in microfluidic devices [Fig2C]. The exceptions to this were the parameter *t*_0_, which was always obtained differently, and *v*_0_, which was pre-determined in the case that a cell cycle occurred because of cell division at the end of a previous simulated cycle. **B**. Generating the initial simulated cell cycles corresponding to synchronised daughters obtained by elutriation. First, parameter sets for many daughter cell cycles were generated (excluding parameter *t*_0_). Then we assigned *t*_0_ values for these cycles such that the cell volume distribution at simulation time *T*_0_ was the same as the one measured experimentally immediately following synchronisation. **C**. Cell division was simulated whenever a cell cycle reached cytokinesis prior to the simulation end. A new mother cycle (1) and a new daughter cycle (2) were generated, both with *t*_0_ set to the moment of cytokinesis of the preceding cycle. The mother *v*_0_ was set as the volume at budding in the previous cycle (blue arrow and lines). The daughter *v*_0_ was set as the total volume grown after budding in the preceding cycle (green arrow and lines). The remaining parameters were randomly sampled for both cycles independently.

To initialise a simulated population, we generated a set of daughter cells whose volume distribution matched the corresponding experimental population immediately after elutriation [Fig. 3B]. We defined the simulation start time, designated *T*_0_, as corresponding to the completion of elutriation in the associated experimental time course. There were two problems to solve in generating this initial population: first, we did not know the cell’s birth times, *t*_0_. Second, the cell volumes at *T*_0_ had to match the experimental volume distribution, whereas the cells’ volumes were dependent on the cycles’ *t*_0_ values. We solved both problems simultaneously with a rejection sampling strategy, where we made use of the fact that we could calculate the volume of a cell at any point in its cell cycle duration from its initial volume, volume growth rates, and cell cycle phase durations. Essentially, we sampled sets of parameters for many daughter cell cycles and, for those cells which could fit within the required volume distribution at some point in their cell cycle, we selected *t*_0_ values prior to simulation start which led to the population achieving the required volume distribution at *T*_0_.

To simulate the subsequent growth and division of the population, whenever a cell divided during the simulated multi-hour time course, we generated a new mother cell cycle and added a new daughter cell to the population. There were two main actions to perform when a cell cycle reached cytokinesis [Fig. 3C]. First, we generated the new mother cycle, set its initial time (*t*_0_) to the point of cytokinesis of the previous cycle, set *v*_0_ as the volume the cell previously had at budding, and sampled the remaining cell cycle parameters from multivariate distribution M. Second, we created a new daughter cell with the same *t*_0_, set *v*_0_ as the total volume grown after budding in the previous cycle, and sampled the remaining cell cycle parameters from multivariate distribution D. We repeated this process of adding subsequent mother and daughter cycles whenever a cell cycle reached cytokinesis prior to the end of the simulated experiment. Thus, by setting up a modelled population in this manner, we could simulate an elutriated population of cells from elutriation up to the end of a several-hour time course experiment.

### Population model fine-tuning and convolution matrix calculation

To test the fidelity of the predictions of our model, we compared simulation results with experimental data from three elutriated populations. In these experiments, we had determined at several time points the cell cycle phase distributions with microscopy using fixed cell samples and the cell volume distributions with a cell counter. We found the largest discrepancy between simulation and experiment in the cell cycle phase distributions, with simulated cells reaching START roughly 20-50 min earlier than in the experiment [Fig. 4A, compare Standard model with Measured]. By contrast, the volume distributions matched reasonably well, although the simulations lacked a well-defined secondary peak at ∼20 fl, visible in the experimental data from around 180 min after elutriation [Fig. 4B, compare Standard model with Measured]. We conjectured that the disagreement in cell cycle phase progression was due to stress caused by elutriation or differences in growth between shake flasks and microfluidic devices, potentially leading to a prolonged initial G1 phase in experimental populations that the model parameters did not account for.

**Figure 4.**
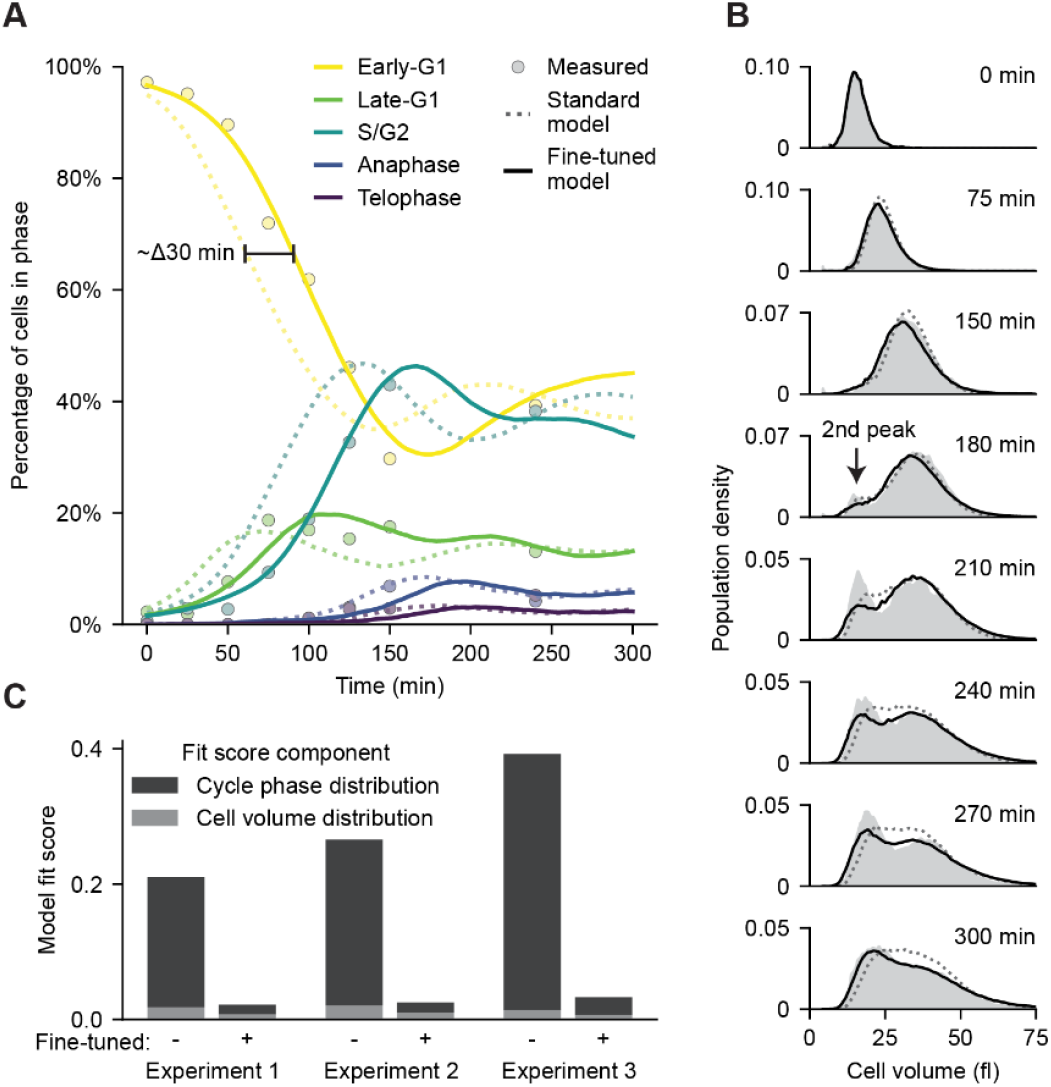
Fine-tuning log-normal cell cycle parameter distributions enabled accurate simulation of experimental populations. **A**. Comparison of cell cycle phase distributions for one time course experiment (Measured), and two corresponding simulated populations generated either with parameter distributions as in Fig. 2 (Standard model) or parameter distributions fine-tuned to the specific experiment (Fine-tuned model). Comparisons for all three experiments are in Supp. Fig. 2 A, C, & E. **B**. Comparison as in **A** for cell volume distributions at selected time points. Comparisons for all time points and all three experiments are in Supp. Fig 2 B, D, & F. **C**. The model fit score was the sum of two components: the sum of squared differences between simulated and experimental cell cycle phase distributions, and the sum of squared differences between simulated and experimental cell volume distributions. Larger model fit scores indicate greater divergence between modelled and experimental populations. After fine-tuning the cell cycle parameter distributions to specific experiments, the fit scores reduced substantially and the cell cycle phase distribution contribution no longer dominated the overall fit scores to the same extent. Experiments 1, 2, and 3 were biological replicate time course experiments beginning with synchronised populations obtained by centrifugal elutriation.

To improve the agreement between simulated and experimental populations, we performed fine-tuning of the parameters that define the multivariate log-normal cell cycle parameter distributions (i.e. the log-mean and log-standard deviation) for each experiment. We used particle swarm optimisation where we allowed log-mean and log-standard deviation values for all the individual cell cycle parameter log-normal distributions to vary within bounds [Supp. Table 3] around their previously estimated values, while keeping the correlations between log-parameters fixed. The most notable change in the post-fitting log-normal distributions [Supp. Table 4] was that the average daughter early-G1 duration increased for all three experiments. Consequently, the differences in cell cycle phase progression between simulated and experimental populations reduced substantially [Fig. 4A, compare Fine-tuned model with Measured]. This change was reflected in the model fit scores, where the contribution from the cell cycle phase data was less dominant in the total score after fitting [Fig. 4C]. There were also improvements in simulated cell volume distributions as the fine-tuned models displayed two distinct volume peaks which better resembled the ones in the experimental data at later time points [Fig. 4B, compare Fine-tuned model with Measured]. As such, the fine-tuning process enabled our simulated populations to approximate experimental populations much better, as assessed by cell volume distribution and cell cycle phase distribution.

Critically, we could now use a fine-tuned model to estimate the convolution kernel for a partially synchronised yeast population in a time course experiment, in the form of the convolution matrix [Fig. 1D], which was essential to deconvolve population-level protein concentration measurements. To do so, we first had to determine which ranges of *φ* values corresponded to each cell cycle phase, beyond the daughter early-G1 (which was pre-defined as -1–0). To this end, we calculated the mean duration of each of these phases for simulated cells and spread these mean durations proportionally over the START-to-START progression between 0 and 1 [Fig. 5A]. Then, for each simulated cell at each time point, we determined its cell cycle phase, the fraction of that phase that it had completed, mapped the cell to the corresponding *φ* position [Fig. 5B], and calculated its volume. Lastly, we binned cells into 20 bins based on their *φ* positions and calculated the fraction of the total population cell volume in each *φ* bin to obtain the convolution matrix [Fig. 5C]. This approach enabled us to precisely estimate the convolution matrix at sub–cell cycle phase-resolution.

**Figure 5.**
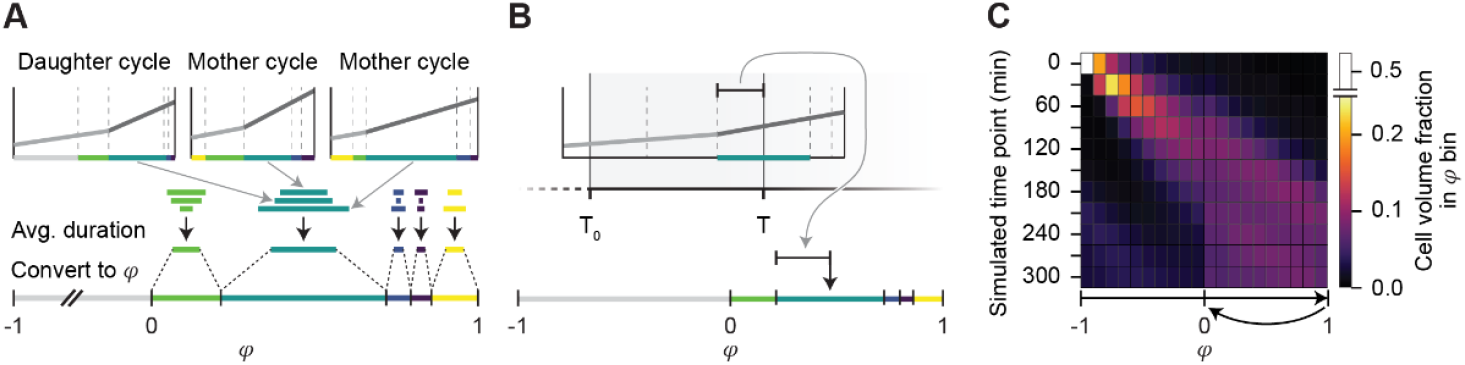
Estimating a convolution matrix for a time course experiment from a corresponding population model. **A**. The ranges of *φ* corresponding to the cell cycle phases late-G1, S/G2, anaphase, telophase, and mother early-G1 were calculated from a fine-tuned population model before estimating the convolution matrix for the corresponding experiment. The phases were distributed over 0 ≤ *φ* ≤ 1 proportional to their mean durations. **B**. At each time point, each cell was mapped to a *φ* coordinate based on its cell cycle progression. The fraction of the current cell cycle phase was found, and the cell was assigned a *φ* coordinate corresponding to the fraction of the region of *φ* corresponding to the cell’s current cycle phase. **C**. A heatmap illustration of the convolution matrix for a single time course experiment obtained from a fine-tuned population model. Each square corresponds to an element of the convolution matrix.

In summary, we developed a computational model capable of simulating the time evolution of the population-level volume and cell cycle phase distributions of an initially partially synchronised yeast culture. Each modelled population consisted of a collection of simulated single cells and their cell cycles. The parameters defining each cell cycle were sampled from multivariate log-normal distributions whose defining parameters were initially based on time-lapse microscopy experiments. The accuracy of a given modelled population was then maximised by fine-tuning the multivariate parameter distribution based on data collected during the corresponding time course experiment. As a result, we could estimate the convolution matrix, which reflected the desynchronisation present within a specific experimental population.

### The regularised least squares deconvolution algorithm

We now required an algorithm that could solve for the mean cell cycle-dependent single cell concentration dynamic of a protein, **c**, given population-level concentration measurements, ^*m*^**c**, and the convolution matrix, **F**. In the ideal case, we could perform deconvolution by directly solving the convolution equation ^*m*^**c** = **Fc** using linear algebra. However, this is an ill-conditioned problem, meaning that even limited noise in the input data can dramatically distort the solution when solving directly or when using simple approaches such as non-negative least squares. This can be seen by taking a known input (such as a sinusoid waveform), generating synthetic ‘measured’ data by convolving the input with a convolution matrix from our elutriation experiments, and comparing the deconvolved results with or without even a small amount of added Gaussian noise [Fig. 6A]. Real datasets contain substantial non-biological variation, originating from instrument noise or from artefacts during sample preparation. Therefore, we required a deconvolution algorithm that found solutions which the fit input data well while remaining robust to noise.

**Figure 6.**
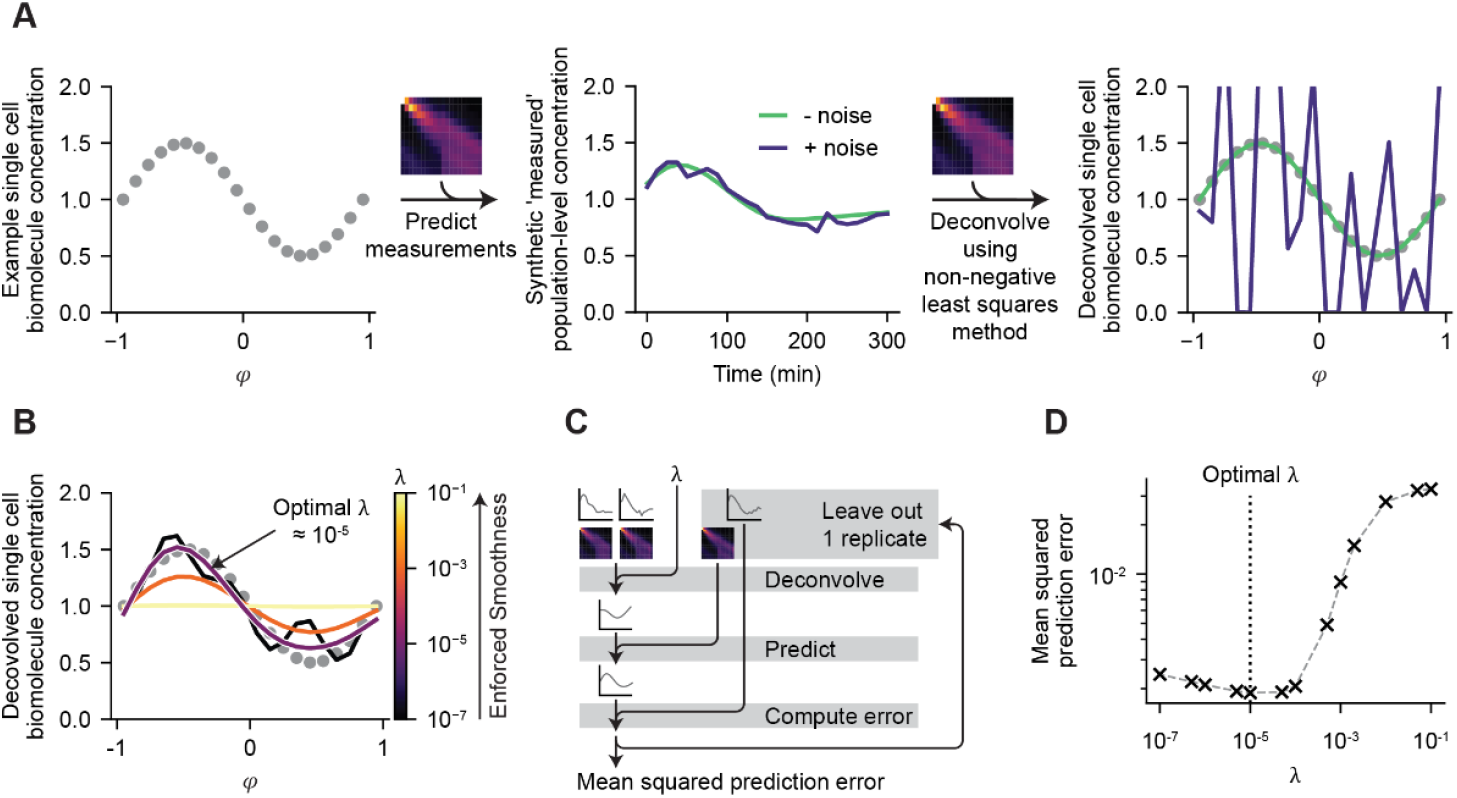
Estimating cell cycle-dependent single cell biomolecule concentrations by regularised deconvolution. **A**. Illustration of the ill-conditioned nature of the deconvolution problem. A known sinusoidal input (left panel) was used to generate synthetic data simulating measured population-level protein concentrations by convolving with a convolution matrix from a fine-tuned population model (middle panel). Performing deconvolution in the ideal no-noise case with a simple non-negative least squares algorithm works perfectly (right panel, compare green line with grey dots). Using the same algorithm on synthetic data with addition of a small amount of Gaussian noise fails with dramatic amplification of noise (right panel, compare purple line with grey dots). **B**. Deconvolution using a regularised least squares algorithm can be more robust to noisy data. Regularisation is implemented as a penalty to the roughness of a given solution scaled by a regularisation parameter, *λ*. Higher values of *λ* enforce greater smoothness of the solution, while lower values permit more dynamic deconvolved solutions. In this specific example with synthetic data, optimal solutions can be found with *λ* ≈ 10^−5^. **C**. A leave-one-out cross-validation approach to determine the optimal *λ* value for deconvolving data for a specific protein. A range of *λ* values were tested between 10^−7^ ≤ *λ* ≤ 10^−1^. **D**. Selection of the optimal *λ* as the one which minimises the mean squared prediction error as obtained in **C**.

In general, we expected cell cycle-dependent protein concentration dynamics to vary relatively smoothly, i.e. without very abrupt concentration changes, and so we selected a regularised least squares algorithm that favours smoother solutions. To this end, we performed deconvolution for a given protein by using an optimisation method to find a solution, **ĉ**, which approximates **c**, by minimising a cost function. This cost function is the sum of two terms: the left-hand term is a standard least squares term which pushes solutions towards close agreement with the experimental data. The right-hand term instead penalises potential solutions based on their roughness, countering the tendency towards extreme output. We calculated the roughness as the sum of squared second order central finite differences of **ĉ** over *φ*. This roughness penalty is scaled by a regularisation coefficient, *λ*, which determines the relative contribution of the two equation terms to the overall cost. A large *λ* value pushes solutions towards increased smoothness (i.e. straight lines), whereas small *λ* values allow for more dynamic solutions [Fig 6B]. The cost function we used had the basic form:

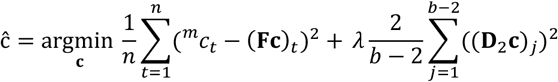

where **D**_2_ is matrix encoding the calculation of second order central finite differences of **c** and 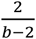 normalises the roughness value based on the number of *φ* bins and the bin width. The exact formulation of the cost function is detailed in Methods section “Deconvolution”, describing additional details to account for normalisation of the input data and the estimation of relative, rather than absolute, concentrations.

The regularisation coefficient *λ* profoundly influences the solution and thus it was important to find appropriate values for it. Ideally, the specific *λ* value would depend on the signal-to-noise ratio in the measured data and might be different for different proteins; for a highly dynamic proteins (high signal-to-noise ratio), a lower *λ* prevents over-smoothing the solution, whereas for non-dynamic proteins (lower signal-to-noise ratio), increased smoothing is reasonable as variation in the measured data would be primarily noise. To identify suitable *λ* values for each protein, we made use of the fact that we had multiple replicate time course measurement series, and therefore we could find reasonable *λ* values using a leave-one-out cross-validation approach. To do this, we performed multiple deconvolutions over a wide range of *λ* values, each time leaving out data from a single replicate experiment. Then we predicted the measured data from the excluded replicate by convolving the solution, obtained by deconvolving the other replicates, with the convolution matrix from the excluded replicate, and compared that prediction with the actual measured data from the excluded replicate [Fig 6C]. We selected the *λ* value which resulted in the lowest mean squared prediction error in these comparisons [Fig 6D]. Using this cross-validation approach, we could select suitable *λ* values for each protein in the dataset.

### Deconvolving time course proteomics experiments

Next, we used our population model and deconvolution approach to estimate cell cycle-resolved protein concentrations from time course proteomics experiments. We grew a pre-culture in glucose minimal medium in a turbidostat, which allowed us to keep cells at high population densities while maintaining an exponential growth rate under stable environmental conditions. We started a time course experiment by separating cells by size using centrifugal elutriation. We eluted the fraction containing the smallest cells, i.e. predominantly early-G1 daughters, and transferred them to a shake flask for further growth. In three replicate time course experiments, we collected three samples every 20 minutes, up to 280 minutes after elutriation: one sample was for proteomics, another to determine the cell cycle phase distribution by fluorescence microscopy, and a third to measure the cell volume distribution [Fig 7A]. We performed global proteome analysis using 16-plex tandem mass tags (TMT) and LC-MS/MS, which provided a dataset allowing comparison of relative protein concentration changes between samples for 3373 proteins.

**Figure 7.**
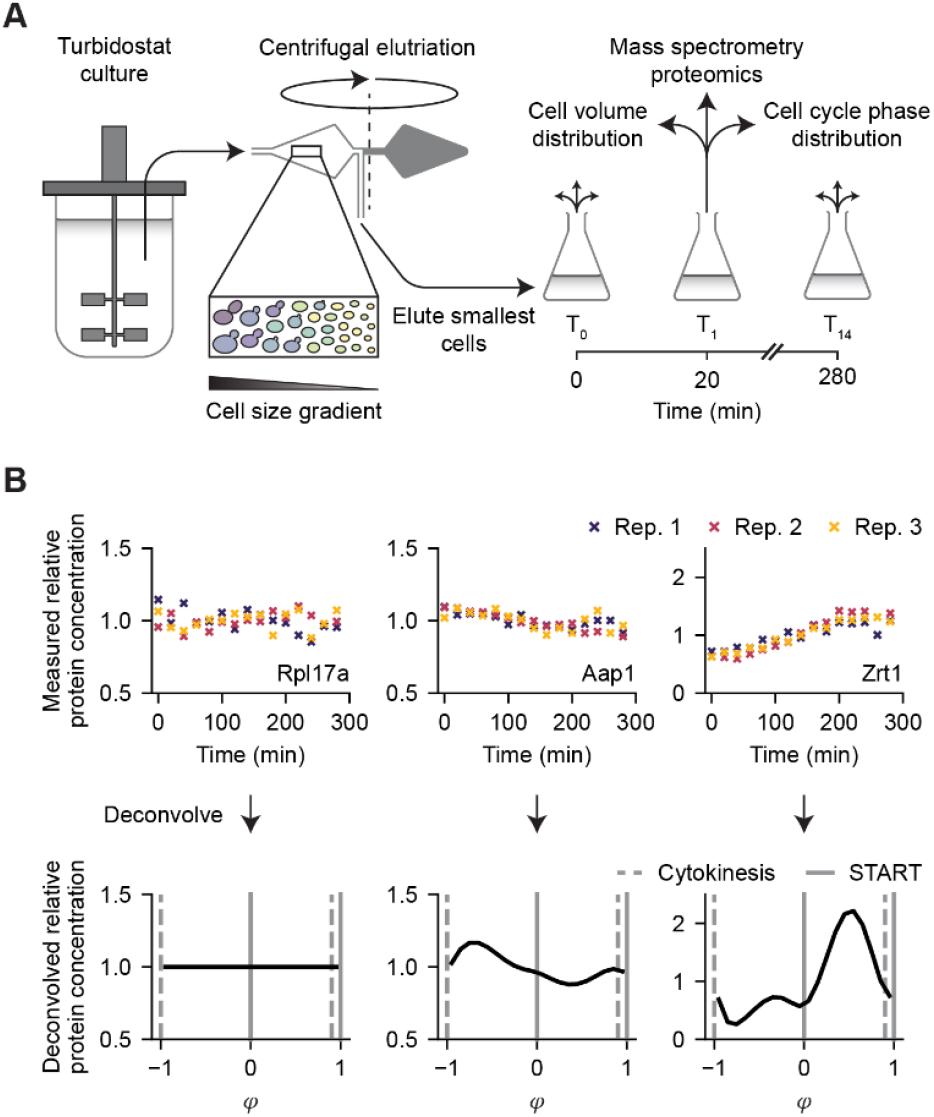
Deconvolution of mass spectrometry proteomics data collected from partially synchronised yeast cultures during time course experiments. **A**. Illustration of the experimental method used to collect samples for population model fine-tuning and for proteomics. The first samples were collected following elutriation and subsequent samples were taken every 20 min up to 280 min inclusive. **B**. Measured and deconvolved relative protein concentrations for three example proteins from three replicate time course experiments. Measured relative protein concentrations were obtained from 16-plex tandem mass tag LC-MS/MS proteomics. Deconvolution, including selection of the regularisation parameter *λ*, was performed separately for each of 3373 proteins using convolution matrices obtained from fine-tuned population models corresponding to the three replicate experimental populations.

We then deconvolved the measured relative concentration data for each protein. First, for each of the three replicate experiments, we fine-tuned a population model using the respective dynamic cell volume and cell cycle phase distribution data [Supp. Fig. 3]. From those models, we then calculated the convolution matrices. When performing the deconvolution, we used our regularised least squares method and imposed the additional constraints that the two values at START (*φ* = 0, 1) should be equal as well as the two values at birth/cytokinesis (*φ* = −1, *φ*_*cyt*_), since those two events each occurred twice on the cell cycle progression axis *φ* [cf. Fig. 1A]. We determined the regularisation factor *λ* for each protein with the outlined cross-validation procedure, wherein we tested *λ* values in the range [10^−7^, 10^−1^] to find the most suitable value [Fig 6C, D]. We expected reasonable *λ* values to lie between those bounds because *λ* < 10^−7^ tended to produce solutions with excessive roughness (i.e. clearly non-biologically realistic solutions) and *λ* = 10^−1^ was already high enough to produce non-dynamic solutions. Overall, we performed deconvolution for the 3373 measured proteins, where we obtained a variety of dynamic and non-dynamic solutions, as indicated with examples [cf. Fig. 7B].

Before performing further downstream analyses with the deconvolved trajectories, we aimed to exclude the solutions with spurious dynamics. False dynamics can occur when an incorrect *λ* value is selected in the cross-validation step, potentially due to chance alignments of errors across different replicates. Aiming to identify the proteins most likely to have been deconvolved correctly, we conjectured that their deconvolved solutions would tend to be more accurate if the respective mass spectrometry data had higher “signal” (i.e. variance due to cell cycle-dependent processes) compared to noise (i.e. variance from error processes). To calculate the “signal” and noise in the measurement data for a given protein, we first estimated the hypothetical noise-free mass spectrometry measurements by re-convolving the solution obtained from deconvolution. We then divided by the mean, centred these values around zero, and calculated the “signal” by summing the squared values. We next calculated the noise as the summed squared residuals between the re-convolved solution and the measurement data and finally obtained the resulting “signal”-to-noise ratio. In addition, since we were most interested in the proteins with the greatest concentration dynamics, we quantified the dynamics of each deconvolved protein trajectory as the peak-to-trough ratio, i.e. the maximum value divided by the minimum value in the deconvolved solution. For downstream analysis, we retained the 563 proteins that were in the upper quartile for both “signal”-to-noise ratio and peak-to-trough ratio, on the basis that those proteins were the ones with the most dynamic trajectories (peak-to-trough ratios ranging approx. 1.3–32) resolved with higher-confidence [Supp. Fig 4].

### Validating deconvolved protein concentration dynamics

To validate the deconvolved trajectories, we tested whether the dynamics of the 563 proteins, in whose deconvolved solutions we had the highest confidence, were consistent with known yeast cell cycle biology. We performed hierarchical clustering of the deconvolved protein concentration trajectories using Pearson correlation as the distance metric and grouped the protein trajectories into five clusters. To test for shared biological functions within each cluster, we performed GO enrichment analysis using the “Biological Process” annotation. In four clusters, we found at least one significantly enriched GO term (adjusted p < 0.05, Fisher’s exact test with Benjamini-Hochberg correction) [Fig. 8B, full GO enrichment analysis results in Supp. Table 5]. The fact that we could identify functional enrichment in proteins with similar cell cycle dynamics lends support to the validity of our deconvolution process.

**Figure 8.**
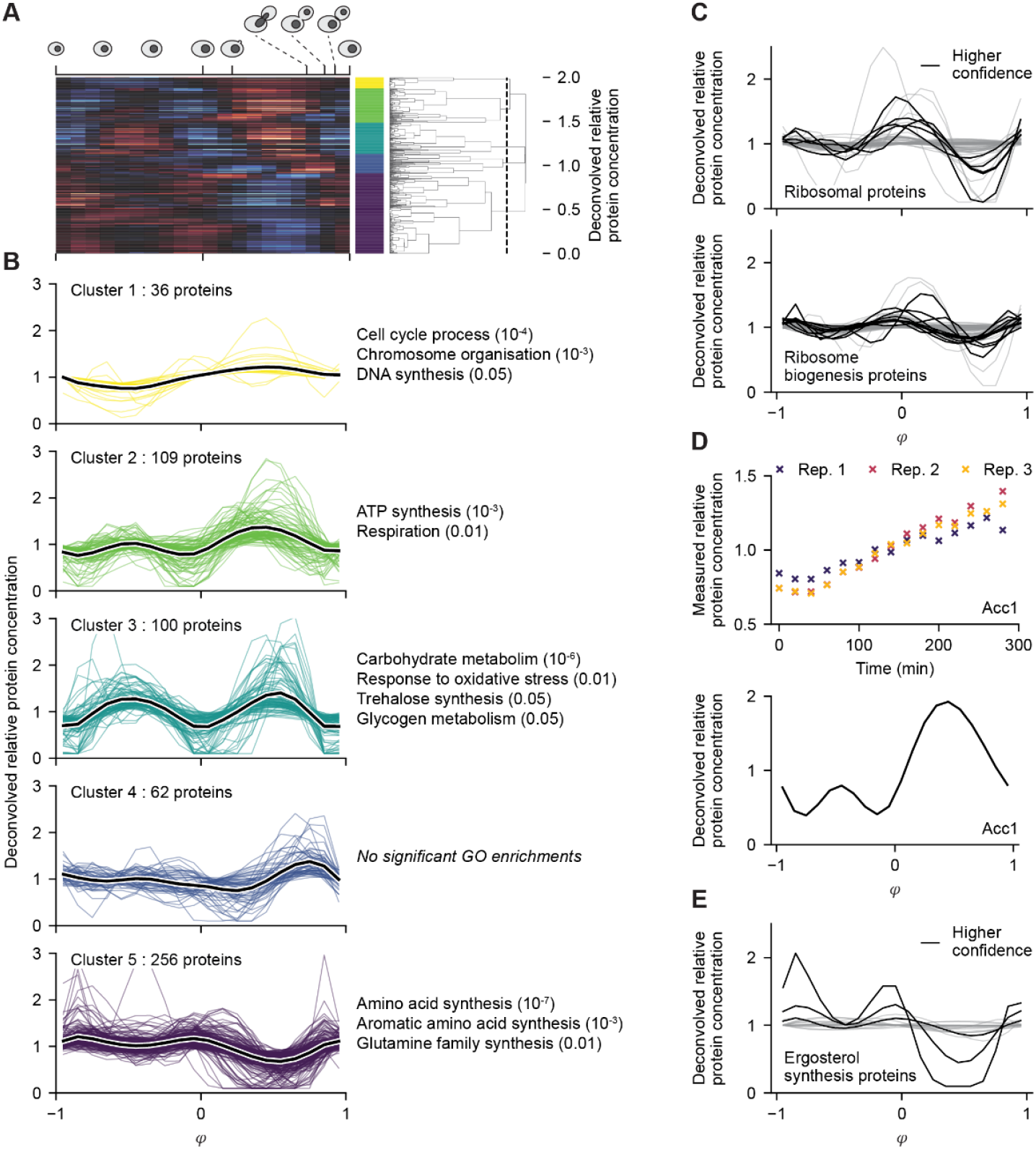
The deconvolved solutions for a higher confidence subset of proteins are consistent with known yeast cell cycle biology. **A**. Hierarchical clustering of the deconvolved solutions for the 563 proteins in our higher-confidence subset of measured proteins. Solutions were clustered based on Pearson correlations using the ‘complete linkage’ method and assigned to one of ten clusters. The higher confidence solutions were selected as those in the top quartile of both estimated signal-to-noise ratio in the measured mass spectrometry data and the top quartile of peak-to-trough ratio in the deconvolved solutions. **B**. Deconvolved solutions for proteins were assigned to one of 5 clusters. For each cluster, coloured lines are the solutions for individual proteins and the black line is the mean across solutions. GO term enrichments were determined using two-sided Fisher’s exact tests with adjustment for multiple testing using the Benjamini-Hochberg method. Labels on the right-hand side of the plots indicate a summary of the enriched GO annotations for each cluster with numbers in parentheses indicating the approximate order of magnitude for the associated adjusted p-values. Full GO enrichment results are in Supp. Table 5. **C**. Deconvolved solutions for cytosolic ribosomal proteins and ribosome biogenesis proteins. Ribosomal proteins (105) were selected as the subset of measured proteins annotated with “Cellular Component” GO term “cytosolic large ribosome subunit” (GO:0022625) and/or “cytosolic small ribosome subunit” (GO:0022627). Ribosome biogenesis proteins (95) were similarly selected as the subset annotated with “Biological Process” annotation “ribosome biogenesis” (GO:0042254), “ribosomal large subunit biogenesis” (GO:0042273), and/or “ribosomal small subunit biogenesis” (GO:0042274). Black lines indicate proteins in the higher-confidence subset (5 for ribosomal proteins, 10 for ribosome biogenesis proteins), grey lines are used for the rest. **D**. Measured and deconvolved data for Acc1, the rate limiting enzyme for fatty acid synthesis from acetyl-CoA. Measured data is from three replicates, the deconvolved solution was obtained by deconvolution of all replicate data together using the regularisation parameter *λ* found by cross-validation. The penultimate timepoint in replicate 2 was excluded for all proteins due to a high proportion of outliers. **E**. Deconvolved solutions for ergosterol synthesis proteins. These were the 21 measured proteins annotated as part of the “superpathway of ergosterol biosynthesis I” on the YeastPathways website. Black lines indicate proteins in the higher-confidence subset (3 proteins), grey lines are used for the rest.

Clusters 1 and 2 both showed lower concentration in daughter early-G1 and higher concentration in S/G2. Cluster 1 was significantly enriched for GO:0022402 “cell cycle process” (cluster mean peak-to-trough ratio = 1.61) [Fig. 8B, Cluster 1]. Importantly, this enrichment was observed despite the absence of yeast cyclins (Cln and Clb), whose dynamic expression is well documented (Wittenberg *et al*, 1990; Ghiara *et al*, 1991; Kelliher *et al*, 2018), but which were not detected in our mass spectrometry data, likely due to their generally low abundance. Higher concentration in S/G2 for this cluster is consistent with enrichments for proteins involved in nucleotide/DNA synthesis and chromosome organization. Cluster 2 proteins had a pronounced peak around S/early-G2 and the cluster was particularly enriched for terms related to aerobic respiration and ATP synthesis (cluster mean peak-to-trough ratio = 1.80) [Fig. 8B, Cluster 2], which is consistent with reported higher O_2_ consumption during S/G2 (Kaspar von Meyenburg, 1969; Takhaveev *et al*, 2023). Overall, both clusters were enriched for proteins involved in processes that are more active during post-budding cell cycle phases.

Clusters 3 and 5 were particularly enriched for metabolic functions, consistent with known patterns of cell cycle-dependent biosynthetic activity. Cluster 3 was enriched for proteins involved in carbohydrate metabolism, which showed concentration peaks during the daughter G1 phase and S/G2 (cluster mean peak-to-trough ratio = 2.07) [Fig. 8B, Cluster 3]. These are both phases that require cell wall synthesis to support growth of either the cell or the bud and a peak in polysaccharide synthesis rate in S/G2 has been reported recently (Takhaveev *et al*, 2023). Cluster 5, where protein concentration peaks around START and drops during S/G2 (cluster mean peak-to-trough ratio = 1.78) [Fig. 8B, Cluster 5], showed strong enrichment for amino acid biosynthesis proteins, which were also found to be enriched among the periodic proteome in another recent study (Campbell *et al*, 2020). Conceivably, a peak in amino acid synthesis around START, in line with increased branched-chain amino acid synthesis as reported for G1 (Blank *et al*, 2023), could contribute to a recently reported rise in protein synthesis around START (Takhaveev *et al*, 2023; Litsios *et al*, 2019). Thus, the enrichments found in these clusters also indicate that deconvolution successfully identified protein concentration dynamics relating to known cell cycle-dependent metabolic changes.

To further compare the deconvolved dynamics with published literature, we considered cytosolic ribosomal proteins and ribosome biogenesis proteins. A high-throughput fluorescence microscopy study using GFP-tagged proteins found a peak in ribosomal proteins and ribosome biogenesis proteins around late-G1 (Litsios *et al*, 2024) while mass spectrometry proteomics results have been more equivocal; one study reported a small G1 peak for both protein groups (Campbell *et al*, 2020) while another argued for no particular cell cycle-dependent dynamics (Blank *et al*, 2020). In our higher confidence subset, there were five (of 105 total) ribosomal proteins and ten (of 95 total) ribosome biogenesis proteins, which all displayed a concentration peak around mid/late-G1 [Fig. 8C]. Conceivably, more data may be needed to reliably identify low amplitude dynamics by deconvolution. Yet, given the consistent G1 position of the peaks identified, along with generally low amplitude dynamics, we interpret our results as being consistent with recent data suggesting a moderate concentration peak somewhere around late-G1 (Litsios *et al*, 2024; Campbell *et al*, 2020), which could functionally underlie a coincident translation rate maximum at this moment in the cell cycle (Takhaveev *et al*, 2023; Litsios *et al*, 2019).

Lastly, we looked for evidence for dynamics within lipid biosynthesis, which did not show specific enrichment in any cluster among our higher-confidence subset, but where several recent reports have identified cell cycle-dependent periodicity. Acc1, the rate-limiting enzyme in fatty acid synthesis (Ivessa *et al*, 1997), was in the 99^th^ percentile for estimated signal-to-noise ratio, the 95^th^ percentile for deconvolved peak-to-trough ratio, and showed clear dynamic behaviour even in the input mass spectrometry measurement [Fig. 8D, upper panel]. The deconvolved Acc1 trajectory showed a low concentration during most of daughter early-G1, but a sharp rise in late-G1 leading to a peak in S/G2 phase [Fig. 8D, lower panel], consistent with Western blot data showing accumulation of Acc1 roughly in daughter late-G1/S (Blank *et al*, 2017). Notably, the peak during S/G2 also coincides with the peak in lipid biosynthetic rate identified for this phase (Takhaveev *et al*, 2023). Ergosterol is another key component of fungal cell membranes and enzymes of the ergosterol synthesis pathway have been reported to have concentration maxima in G1 (Litsios *et al*, 2024). Consistently, three (of 21 total) of these enzymes were in our higher-confidence subset and all displayed peaks around G1 [Fig. 8E].

Overall, we found that the deconvolved concentration trajectories – at least those in our higher confidence subset – clustered into groups with similar biological functions and the cell cycle-dependent concentrations were broadly consistent with known yeast cell cycle biology. Together, these observations support the validity of our higher confidence deconvolved cell cycle-dependent protein concentration trajectories. As such, we provide a dataset with deconvolved cell cycle-resolved concentration dynamics for the 563 proteins in our higher confidence subset as well as putative dynamics for an additional 2800+ yeast proteins [Supp. Table. 6].

### Transcriptional regulation during the cell cycle

After validating the dataset, we set out to identify regulatory transcription factors (TFs) whose changing activity could explain the identified protein concentration dynamics. To this end, we used documented activating and inhibitory TF-gene relationships with DNA-binding evidence from YEASTRACT (Teixeira *et al*, 2023) for genes whose protein were in our higher confidence set of proteins. In brief, we quantified an evidence score for each TF’s cell cycle-dependent activity by measuring the alignment between the deconvolved concentration changes of the higher-confidence proteins and the TF’s expected effects. As each TF has multiple targets, we then aggregated these measurements into a single, cell cycle-resolved dynamic for each TF summarising the evidence for its activity throughout the cell cycle.

Specifically, to estimate TF activity changes, we computed the log_2_ concentration fold change for each protein between each pair of successive *φ* bins. To prevent proteins with the largest concentration amplitudes from dominating the scores, we normalised each protein’s log_2_ fold changes by dividing by that protein’s maximum log_2_ fold change. Next, we encoded the TF-gene interactions in a matrix where a 1 indicated an activating relationship, a -1 indicated a repressive relationship, and 0 indicated either no relationship or ambiguous evidence. We then determined the matrix product between the normalised log_2_ fold changes and the TF-gene interaction matrix [Fig 9A]. To allow comparison between TFs with different numbers of targets, we divided the resulting values for each TF by the number of its targets within the higher confidence subset of proteins. This yielded a number between -1 and 1 for each TF and bin-to-bin transition; a value of 1 indicated that all targets of a TF underwent their maximum bin-to-bin log_2_ fold change in the direction consistent with the TF’s activity. Conversely, a value of -1 indicated that all of a TF’s targets underwent maximal changes opposite to what would be expected if the TF were active. Finally, we determined the TF activity score as a function of the cell cycle by calculating the cumulative sum of the estimated TF activity changes over *φ*. While not a direct measurement of TF activity, we refer to this TF activity score as “TF activity” below for brevity.

**Figure 9.**
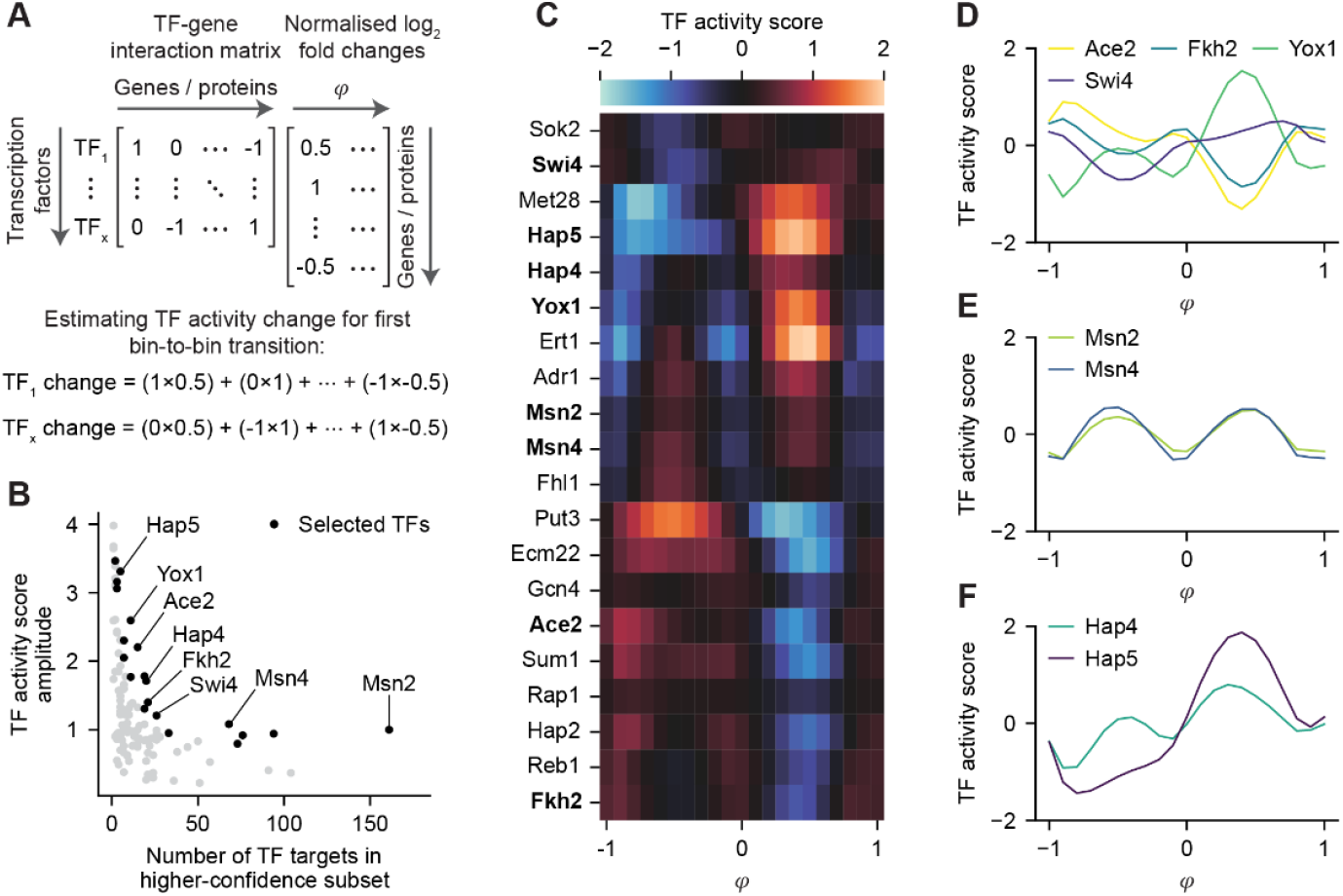
Estimation of cell cycle-dependent transcription factor (TF) activity from deconvolved solutions. **A**. Illustration of a critical step in the estimation of TF activity changes from protein deconvolved solutions. TF-gene relationships with DNA-binding evidence were obtained from YEASTRACT (Teixeira *et al*, 2023) for genes whose protein products were in our higher confidence set of proteins. The TF-gene interaction matrix was constructed such that a 1 indicated an activating relationship, a -1 indicated a repressive relationship, and 0 indicated either no relationship or ambiguous evidence. The matrix of bin-to-bin log_2_ fold changes was obtained by first finding the difference between successive *φ* bins in the log_2_-transformed deconvolved solutions for the higher confidence set of proteins. These differences were then normalised to the range -1–1 by dividing values for each protein by the maximal bin-to-bin log_2_ fold change for the respective protein. The evidence supporting TF activity changes was then quantified as the matrix product of the TF-gene interaction matrix and the normalised bin-to-bin log_2_ fold change matrix. **B**. TF activity scores were calculated as the cumulative sum of TF activity changes over *φ* and centred around the mean value. The TF activity score amplitude roughly summarises the size of inferred cell cycle-dependent TF activity changes. The selected TFs were identified as those with activity score amplitudes in the top 5% compared with 10^5^ trials where an equivalent number of ‘targets’ were sampled randomly without replacement from the set of proteins with higher confidence solutions. TFs indicated by name are those shown in **D, E**, or **F**. **C**. Heatmap showing TF activity scores across *φ* for selected TFs from **B**. TFs with names in bold typeface are those shown in **D, E**, or **F. D, E, F**. Inferred cell cycle-dependent TF activity dynamics for a subset of TFs.

From the 115 TFs in the YEASTRACT dataset with targets in our higher-confidence subset, we aimed to identify those with evidence of substantial dynamic activity during the cell cycle. We based our selection on the amplitude of a TF’s activity; high amplitudes reflect the degree of co-regulation of the TF’s targets over the cell cycle as well as more dynamic protein expression changes. We assumed that, if a TF did not exert cell cycle-dependent regulation, the expression of its targets would tend to be uncorrelated with the known TF-gene relationships. Hence, the contributions from each target to the TF activity would roughly cancel out and result in a low TF activity amplitude. However, this assumption was less valid for TFs with fewer targets since they had a higher chance for most of their targets to be co-expressed, even if the TF did not have cell cycle-dependent activity. This fact resulted in a roughly inverse relationship between TF activity amplitude and the number of TF targets [Fig. 9B]. As such, to estimate the probability of observing each TF’s activity amplitude given its number of targets, we randomly sampled different sets of targets for each TF and calculated the resulting activity amplitudes. For further analysis, we retained those 20 TFs whose measured activity amplitudes were within the top 5% of the values calculated from 10^5^ random samples [Fig 9B] showing different cell cycle-dependent activity profiles [Fig. 9C].

Some of these 20 TFs have known roles in cell cycle-dependent regulation. For example, we found the TF activity of Ace2 to peak around M phase and particularly in daughter early-G1 [Fig 9D]. Indeed, Ace2 is specifically partitioned into the daughter cell nucleus where it promotes destruction of the mother/daughter septum and contributes to the longer daughter early-G1 (Laabs *et al*, 2003; Di Talia *et al*, 2009; Colman-Lerner *et al*, 2001). Another TF, Yox1, represses G1 and M phase genes during S/G2 (Pramila *et al*, 2002), which is precisely where we inferred the highest TF activity. Similarly, the forkhead transcription factors Fkh1 and Fkh2 act on a cluster of genes in late-G2/M phase (Kumar *et al*, 2000; Zhu *et al*, 2000; Koranda *et al*, 2000), which is the point at which we see an increase Fkh2 activity. On the other hand, inferred Swi4 activity slowly increased from mid-G1 towards M phase, which was unexpected given that it acts in the SBF complex to activate genes involved in the G1/S transition and is later repressed (Amon *et al*, 1993; Koch *et al*, 1993; Skotheim *et al*, 2008). However, only 26 of the 162 Swi4 targets with unambiguous directional regulation and DNA binding evidence in the YEASTRACT database were in our higher confidence subset of proteins, which may have been insufficient to give a robust readout of canonical Swi4 activity. Taken together, the activity profiles for multiple TFs closely aligned with their known behaviour, which validates our approach for identifying potentially unknown cell cycle-dependent regulators.

This analysis allowed us to discover new cell cycle-dependent TF activities. For instance, we found that the activity for Msn2 and Msn4, two key TFs downstream of PKA and TOR (Görner *et al*, 1998; Beck & Hall, 1999), showed a pronounced peak in daughter early-G1 and in S/G2 [Fig 9E]. While activity of Msn2/4 is typically associated with diverse stress responses (Martínez-Pastor *et al*, 1996; Gasch *et al*, 2000), there are also indications that they may have cell cycle-dependent activity. For example, transcriptomics analysis identified Msn2 targets as having high periodicity consistent with Msn2 activity in early-G1 (Teufel *et al*, 2019). Furthermore, Msn2/4 promote the acetyl-CoA accumulation in synchronously dividing chemostat populations during the prolonged quiescent phase between division cycles (Kuang *et al*, 2017). As such, we suggest that Msn2/4 may promote similar metabolic conditions in daughter cells during their extended early-G1 phase. At present, it is less clear what role Msn2/4 may play during S/G2. However, it is notable that Msn2/4 TF activity score dynamics occur in anti-phase to cell cycle-dependent oscillations in TORC1 and PKA activity (Guerra *et al*, 2022), both of which are negative regulators of Msn2/4. Given that TORC1 and PKA are central metabolic regulators, it is conceivable that part of their downstream action may be mediated via rhythmic repression and de-repression of TFs like Msn2/4, potentially helping to orchestrate the differential timing of cell cycle biosynthetic activity (Takhaveev *et al*, 2023).

We also identified TFs that could contribute to cell cycle-dependent switching between fermentative or respiratory metabolic regimes. Yeast grown in high-glucose conditions show maximal glucose update rates and glycolytic flux in G1 (Takhaveev *et al*, 2023; Monteiro *et al*, 2019). Furthermore, the G1/S transition is marked by an increase in oxygen uptake, CO_2_ excretion and a decrease in ethanol production (Takhaveev *et al*, 2023; Kaspar von Meyenburg, 1969) which reverses around the beginning of M phase – suggestive of a rebalancing from fermentative to respiratory growth during S/G2. Interestingly, we found Hap4 and Hap5 (though not Hap2) activity scores to peak around S/G2, which is intriguing given that Hap2/3/4/5 complex activity generally promotes mitochondrial function and respiratory metabolism (Lascaris *et al*, 2002; Buschlen *et al*, 2003). As such these TFs may contribute to the metabolic switching during the cell cycle as identified previously.

In summary, the results of our TF activity analysis reinforce the capacity for our deconvolution method to reveal both known and potentially unknown cell cycle-dependent biology. Using the calculated TF activities, we identified several TFs with known matching cell cycle-dependent behaviours. Furthermore, we propose potential roles for regulators like Msn2/4 and the Hap2/3/4/5 complex in shaping metabolic dynamics to meet changing demands during a full division cycle.

## Discussion and conclusion

In this paper, we resolved budding yeast cell cycle-dependent protein concentration trajectories by deconvolution of proteomics data. To do so, we developed a computational model, parameterised with data from microscopy experiments, to simulate the time evolution of the population-level volume and cell cycle phase distributions of an initially partially synchronised yeast culture. Then, building on this model, we developed an approach to deconvolve the dynamic concentration trajectories of proteins during the budding yeast cell cycle. Applying this approach to the proteomics data we generated, we estimated the cell cycle-dependent concentration trajectories for 3373 proteins and identified a subset of 563 proteins with higher-confidence cell cycle-dependent dynamics, with peak-to-trough ratios ranging from 1.30 to about 32. We anticipate this dataset will be an important resource for cell cycle research in budding yeast.

After a series of studies that applied deconvolution to transcriptome data in budding yeast, (Bar-Joseph *et al*, 2004; Qiu *et al*, 2006; Rowicka *et al*, 2007; Orlando *et al*, 2007, 2009; Guo *et al*, 2013), to our knowledge, we are the first to quantitatively deconvolve cell cycle-dependent proteomics data in this organism. Deconvolving proteome and transcriptome data presents different challenges; changes in protein abundance are generally lower compared with mRNAs, for instance following environmental stimuli (Lahtvee *et al*, 2017) or during the cell cycle (Campbell *et al*, 2020; Litsios *et al*, 2019), meaning that the dynamics we sought to determine here are generally of lower amplitude and thus harder to isolate. Furthermore, while resolving lower amplitude dynamics from noise is already challenging, the ill-conditioned nature of the deconvolution problem tends to greatly compound the effects of even small errors in the input measured data. Consequently, we had to mitigate the impact of measurement errors by (i) maximising the accuracy of the estimated convolution matrix, (ii) identifying the most suitable regularisation parameter value for each protein individually via cross-validation, and (iii) using conservative filtering such that we retained only the higher confidence deconvolved solutions for downstream analysis.

Our model-based method for estimating the convolution matrix, and the way in which we defined our standardised cell cycle coordinate, offered several advantages. First, we estimated a time-varying convolution kernel for each experimental replicate, which was necessary given that synchrony decreased over (and differed between) our time courses. This contrasts with the use of a time-invariant convolution kernel to deconvolve transcriptomics from cycling chemostat cultures (Rowicka *et al*, 2007). Second, our use of rich single-cell microfluidics data to parameterise our model enabled us to simulate the cycle-to-cycle variation, in terms of individual cell cycle phase durations, volume growth rates, and parameter correlations present in experimental cultures. Furthermore, we fine-tuned each simulated population to match the sampled culture using cell cycle progression data and cell volume distributions collected from the same culture used for proteomics. In contrast, previous models used to deconvolve transcriptome data typically either used budding index data and/or DNA content measurements from flow cytometry (Orlando *et al*, 2009; Guo *et al*, 2013; Orlando *et al*, 2007; Bar-Joseph *et al*, 2004), which provides less information about cell cycle phase distribution in G1 and M phases. Finally, with our approach, we could deconvolve protein concentration dynamics separately for daughter and mother early-G1 phases, which was also a feature of the CLOCCS models used by Haase and Hartemink for deconvolution of transcriptome data (Orlando *et al*, 2009, 2007; Guo *et al*, 2013).

Nonetheless, perhaps the most substantial improvement toward estimation of the convolution matrix was to include cell volume information. The blurring effect of desynchronisation on measured values arises from heterogeneity in cell cycle phase progression, changes in cell volume, and asymmetric division. In essence, the process of measuring the average concentration of a biomolecule involves collecting a sample of cells, lysing them, assaying the abundance of each biomolecule of interest, and dividing by the total sampled cell volume. As such, the average concentration is the mean of the concentrations within individual cells weighted by their respective cell volumes. While work in other organisms has attempted to account for cell volumes, such as for the bacterium *Caulobacter* (Siegal-Gaskins *et al*, 2009), prior deconvolution studies in budding yeast all focused exclusively on the cell cycle progression aspect of desynchronisation. However, given the substantial cell volume variation within yeast populations [cf. Fig. 4B, Supp. Fig. 2 & 3 B, D, E], volume is clearly an important factor to incorporate into the convolution kernel. To do so, we explicitly modelled cell volume growth in G1 and S/G2/M separately and propagated volumes to subsequent mother and daughter cell cycles after each cell division. Having simulated both cell cycle progression and cell volume growth in our model, the resulting convolution matrices enabled reconstruction of cell cycle-resolved protein concentration dynamics from bulk proteomic measurements.

Despite the strengths of our approach, it also involves limitations and trade-offs. Due to the ill-conditioned nature of deconvolution processes, any noise in the experimental data poses a particularly difficult challenge. To mitigate this, we used periodicity and smoothness assumptions in our regularised least squares approach to stabilise the inference of protein dynamics. However, these assumptions may not be valid for particularly short-lived and/or dynamic proteins, and so our analysis would likely have underestimated their true dynamic range. Additionally, the selection of a suitable regularisation coefficient *λ* for each protein was vital for an accurate deconvolved solution. While our cross-validation method for *λ* selection helps buffer against noise in individual replicates, excessive variation in the input data, or chance alignments of errors across replicates, can still lead to sub-optimal solutions. Looking ahead, we anticipate future deconvolution methods could benefit from sharing information across multiple proteins to better distinguish signal from noise, rather than deconvolving each protein individually, and/or by using regularisation methods with different characteristics, such as the wavelet-basis approach used previously in a transcriptomics study (Guo *et al*, 2013). Lastly, as always, the success of deconvolution analyses critically depends on high quality experimental data obtained by maximising population synchrony, minimising measurement noise, and dense temporal sampling from several replicates.

Several previous studies have measured the cell cycle resolved proteome (Flory *et al*, 2006; Zhang *et al*, 2019; Blank *et al*, 2020; Campbell *et al*, 2020; Litsios *et al*, 2024) and it is important to interpret those results with the problem of desynchronisation clearly in mind (cf. Fig. 6A left and centre panel). Of the mass spectrometry studies, none applied deconvolution to the generated datasets and, as such, they will have necessarily underestimated dynamics – particularly at cell cycle phases further away from the synchronisation point. Consequently, one must be very cautious about concluding that a protein is not dynamic based on an apparently ‘flat’ trajectory. Nonetheless, given this limitation, it is striking that two papers identified hundreds (Blank *et al*, 2020) to nearly a thousand proteins (Campbell *et al*, 2020) with significant cell cycle-dependent dynamics. These numbers are in the same range as our own total of 563 higher confidence deconvolved dynamic proteins and the ∼800 proteins identified with cell cycle-dependent concentrations in a recent high-throughput microscopy study (Litsios *et al*, 2024). In addition to the identification of dynamic proteins, we advanced upon these previous attempts by reconstructing continuous pseudo-time concentration trajectories which substantially improve on the cell cycle resolution of previously known protein dynamics. Overall, as we get better at detecting low abundance proteins and deconvolving our measurements, the number of proteins with cell cycle-dependent dynamics will likely increase.

We envision that our cell cycle resolved proteome dataset will serve as a valuable resource for future studies of the dynamic processes that coordinate cell growth with division. Given the recent discoveries of autonomous oscillations in metabolism and biosynthesis (Takhaveev *et al*, 2023), their possible origin as a solution to proteome constraints (Wongprommoon *et al*, 2024), and their interplay with cell cycle control (Litsios *et al*, 2019), our data should help uncover relevant protein factors and potential targets for further investigating these phenomena.

## Materials and methods

### Strains

We used a YSBN6 *WHI5*-*mCherry*-*Ble HTA2*-*sfGFP*-*Kan*MX strain for all experiments (Litsios *et al*, 2022). The parental prototrophic MATa haploid YSBN6 (FY3 *ho*::*Hph*MX4) strain (Canelas *et al*, 2010) is derived from the S288c genetic background (Winston *et al*, 1995). We maintained frozen glycerol stocks at -70 °C. To retrieve cells from storage, we streaked a sample onto 2% glucose YPD agar plates and grew cells for 2-3 days at 30 °C. We kept colonies on plates at 4 °C for up to one month.

### Liquid media

We used synthetic minimal media for growth in liquid culture.

For the microfluidics experiments used to determine parameters for the cell population model, as well as the time course experiments used to test the population model, we used 1x YNB without amino acids (Formedium) with 2% glucose (w/v) and 100 mM potassium hydrogen phthalate/potassium hydroxide buffer (pH 5.55), hereafter called “buffered YNB medium”.

For the time course proteomics experiments, we used the medium described in (Verduyn *et al*, 1992) with 10 mM potassium hydrogen phthalate/potassium hydroxide buffer (pH 5.0), hereafter called “Verduyn medium”. For preculture and turbidostat growth, we used Verduyn medium with 2% glucose (w/v). For growth of cells following elutriation, we used Verduyn medium with 1.6% glucose (w/v) and 0.14% ethanol (v/v) to more closely resemble the conditions present in the turbidostat.

### Turbidostat culture

For the elutriation experiments, we needed high-density cultures growing at a high growth rate and under constant environmental conditions. To accomplish this, we grew cultures using a turbidostat bioreactor setup that consisted of a 3 l bioreactor (Applikon) connected to a 20 l medium reservoir, both of which were sterilised before use by autoclaving. During operation, we grew cells in 2 l medium in the bioreactor and maintained the medium temperature at 30 °C using a heating element connected to a water bath. We also constantly bubbled filtered air through the medium (1–2 l min^-1^) to maintain aeration and stirred at 300 rpm with a Rushton/marine impeller.

To allow for the turbidostat operation of the reactor, we built a turbidostat control circuit based on published specifications and software (Hoffmann *et al*, 2017). The control circuit consisted of an OD sensor, a connected computer, and a pair of pumps controlled by an Arduino Nano microcontroller. We mounted the OD sensor around a bulb in a glass tube, through which we pumped bioreactor medium, which allowed for constant tracking of culture density. OD measurements were passed to the connected computer that, after we had entered the desired OD set point, activated one of the peristaltic pumps, via the microcontroller, to inject fresh medium as required to maintain the population density. The other peristaltic pump continually removed medium when the liquid level exceeded 2 l to ensure a constant volume of medium in the bioreactor. The calibration of the turbidostat OD sensor varied between experiments, so we determined the correct set point each time by taking OD_600_ measurements using a spectrophotometer (Amersham Novaspec Plus) or cell count measurements with a CASY (Model TT, Innovatis) instrument.

To prepare cells for turbidostat growth, we first inoculated a 10 ml liquid culture in a 100 ml flask with a single colony from an agar plate and grew cells overnight in a shaking incubator (300 rpm, 30 °C). We diluted cells the next morning into either 25 ml fresh pre-warmed medium in a 250 ml flask or 50 ml medium in 500 ml flask. In the evening, we again diluted cells either into 250 ml fresh pre-warmed medium in a 2 l flask or into 500 ml medium which was split equally across 4x 500 ml flasks to ensure adequate aeration. At this time, we also pumped either 1.75 or 1.5 l fresh pre-warmed medium into the bioreactor, activated the heating circuit, aeration, and the stirrer and allowed the medium to equilibrate overnight. The next morning, we inoculated the bioreactor by transferring the overnight culture into the bioreactor through a small port using a 50 ml pipette. Up to that point, we maintained the cultures at an OD_600_ ≤ 2.

Following inoculation, we allowed the culture to reach the desired cell population density in the bioreactor, which took 2–8 hours depending on the densities of the inoculum and desired turbidostat culture. Once the population density set point was reached, we activated the turbidostat control circuit and allowed the culture to equilibrate for at least 16 h to achieve a steady state prior to use in elutriation experiments. For the time course proteomics experiments, we used a set point corresponding to OD_600_ = 2 as measured with the spectrophotometer. For the microfluidics experiments and time course experiments used to test the population model, we used a set point corresponding to a population density of 20–25 million cells ml^-1^ as measured with the CASY.

### Elutriation

To synchronise cells by centrifugal elutriation, we used an Avanti J-20XPI Centrifuge (Beckman Coulter) equipped with a JE-5.0 elutriation rotor (Beckman Coulter, 356900) and a 40 ml elutriation chamber (Beckman Coulter, 356940). We used a peristaltic pump (Watson-Marlow 520S) to pump culture or medium into the elutriation apparatus and used a three-valve switch (Smiths Medical, model MX2311L) to change between the two. Prior to performing elutriation, we heated the centrifuge to 30 °C and filled the elutriation chamber with fresh pre-warmed medium. We performed all elutriation steps with the centrifuge spinning at 3200 rpm.

For the proteomics time course experiments, we pumped 1 l of turbidostat culture through the elutriation apparatus with a peristaltic pump rate of 8 rpm (flow rate approx. 35 ml min^-1^) to load cells into the elutriation chamber. After loading was completed, we switched from pumping cell culture to pumping fresh pre-warmed elution medium (Verduyn medium with 1.6% glucose (w/v) and 0.14% ethanol (v/v)) at the same flow rate. We then slowly increased the pump rate until the smallest cells in the elutriation chamber, which were visible through the centrifuge viewing port as the so-called “cell boundary” (see Beckman-Coulter manual), approached the exit of the chamber when the pump rate was about 13 rpm (approx. 55 ml min^-1^). Then we increased the pump rate by 1.5 rpm to elute the smallest cells (final flow rate approx. 60– 65 ml min^-1^). We collected 300 ml of outflow in a pre-warmed 1 l flask with a cell density of approximately 4–11×10^6^ cells ml^-1^.

For the microfluidics experiments and time course experiments used to test the population model, we used a slightly different protocol. We loaded the elutriation chamber for 20 min at a pump rate of 11 rpm (approx. 45 ml min^-1^). We switched to pumping fresh pre-warmed medium at the same flow rate for about 1 min and then eluted cells at a pump rate of 14 rpm (approx. 60 ml min^-1^). We collected 300 ml of outflow in a pre-warmed 1 l flask with a cell density of approximately 2–9×10^6^ cells ml^-1^.

### Time lapse fluorescence microscopy of cells

Prior to imaging, we grew cells in a turbidostat culture and collected an early-G1 culture by elutriation. After completion of elutriation, a sample of elutriated cells was loaded onto a microfluidic device with a design as reported previously (Huberts *et al*, 2013). We maintained cells in the microfluidic device at 30 °C using a microscope incubator (Life Imaging Services). We constantly supplied fresh medium (see below) using a syringe pump. We pumped medium at a flow rate of 0.5 µl min^-1^ during the process of loading cells onto the device rate and 3.2 or 4 µl min^-1^ during time lapse imaging.

We imaged cells in the microfluidic device using a Nikon Eclipse Ti-E widefield inverted fluorescence microscope fitted with using an Andor DU-897EX camera. We used a 100x Nikon CFI Plan Apo λD Oil (NA = 1.45) objective and a Lumencor AURA excitation source. For brightfield imaging, we illuminated cells with a halogen lamp, fitted with a 420 nm beam splitter to filter out UV wavelengths, with a 200 ms exposure. To image sfGFP, we excited samples at 470 nm using a 470/40 nm band-pass filter, a 495 nm beam splitter, and a 525/50 nm excitation filter at a light source intensity of 3% over a 200 ms exposure. To image mCherry, we excited samples at 565 nm using a 560/40 nm band-pass filter, a 585 nm beam splitter, and a 630/75 nm emission filter at a light source intensity of 12% over a 300 ms exposure. We operated the microscope using Nikon NIS-Elements software and used the Nikon Perfect Focus System to avoid loss of focus during time lapse imaging. We acquired images in a non-overlapping grid of positions. We started imaging approximately 40–45 min following completion of elutriation and imaged brightfield, sfGFP and mCherry every 5 min for 16 h.

In total, we performed three imaging experiments: in two experiments, cells were cultured in buffered YNB medium. In the third experiment, we again pre-cultured cells in the turbidostat in buffered YNB medium but cultivated the elutriated cells on the microfluidic device in conditioned buffered YNB. We prepared this conditioned buffered medium by pumping 800 ml culture out from the turbidostat, once the population density had reached the target of 25 million cells ml^-1^, and filtered it through a filter with 0.2 micron pore size (Fisherbrand, FB12566510). We did not see substantial differences during the analysis of cells grown in the two different media and so we pooled data from the three experiments during downstream analysis.

### Calculation of cell cycle parameter distributions

We analysed the time lapse image series obtained from cells grown on microfluidic devices to estimate cell volumes and to identify when key cell cycle events occurred. We only analysed cell cycles for cells which were born on the microfluidic device and where the cycle began (i.e. the cytokinesis event preceding the cell cycle) occurred at least 2 h after the start of imaging. We segmented cells and fitted ellipses to their cross-sectional areas in a semi-automated manner using the BudJ plugin (Ferrezuelo *et al*, 2012) for FijiJ/ImageJ (Schindelin *et al*, 2012; Schneider *et al*, 2012). For each tracked cell cycle, we segmented the mother cell as well as the growing bud once it was large enough to be reliably detected by BudJ. We estimated the volume of the mother and bud from the major and minor radii of the fitted ellipses using the assumption that cells had a prolate spheroid shape. To estimate volumes for the bud prior to the point at which we could first track it, we set the volume at the time point when budding occurred to 0 fl and then linearly interpolated between budding and the first tracked time point. We calculated the G1 linear volume growth rate as the gradient of the line between the cell volume at the beginning of the cycle and the volume at the point of budding. We calculated the S/G2/M linear volume growth rate as the gradient of the line between the volume at budding and the summed mother cell and bud volume at the subsequent cytokinesis.

We manually determined the timepoints of the cell cycle events START, budding, karyokinesis, mitotic exit, cytokinesis and separation as follows: START was the first frame where an unbudded cell no longer had a nuclear Whi5-mCherry focus. Budding was the first frame where a post-START cell displayed a dark focus on its surface which subsequently grew into a bud. Karyokinesis was the first frame when the Hta2-sgGFP focus of a budded cell either separated into two, or when it became elongated or two-lobed preceding full separation. Mitotic exit was the first frame when a post-karyokinesis cell displayed Whi5-mCherry foci in both mother and daughter nuclei. Cytokinesis was the first frame after mitotic exit where a clear darkened septum was visible between mother and bud — typically this was the frame after mitotic exit occurred. Separation was the first frame when the mother and daughter cells became separate particles, usually indicated by a sudden increase in distance between the two. We then calculated the durations of early-G1, late-G1, S/G2, anaphase, telophase, and separation for each cycle as the time differences between the relevant cell cycle events [Fig. 2B].

We classified analysed cell cycles as either daughter or mother cycles. A daughter cycle was the first full cycle for a cell following separation from its mother. Mother cycles were any cycle for a cell after its daughter cycle. We excluded any cycle with any negative or zero durations or growth rates. We also excluded as an outlier any cycle where the log of one or more parameter values was more than three standard deviations away from the mean log value for the respective parameter. In total we used data from 192 cycles (134 daughters, 58 mothers) after excluding 5 cycles with negative or zero parameter values (1 daughter, 4 mothers) and 14 outliers (5 daughters, 9 mothers).

To facilitate sampling cell cycle parameters when modelling the growth of yeast populations, we fitted log-normal distributions to the initial volume, both linear growth rates, and all cycle phase durations (including separation) for daughter cycles as well as to the mother cycle early-G1 duration. We combined these into two different multivariate log-normal distributions labelled D and M. Distribution D comprised the log-normal distributions for initial volume, both linear growth rates, all cell cycle phase durations (including separation) for daughter cycles. Distribution M comprised the log-normal distribution for early-G1 for mother cycles, and the log-normal distributions for both linear growth rates, late-G1 duration, S/G2 duration, anaphase duration, telophase duration, and separation for daughter cycles. For both multivariate log-normal distributions, we calculated the covariance matrix from the matrix of Pearson correlations between log-transformed daughter cell cycle values. Specifically, we obtained a covariance matrix **A** ∈ ℝ^*x*×*x*^ from correlation matrix **B** ∈ ℝ^*x*×*x*^ and standard deviation vector ***σ*** as:

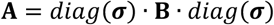

### Time course sampling from elutriated cultures

We performed post-elutriation time course sampling experiments to (i) collect data to test the yeast population model and to (ii) perform mass spectrometry proteomics. In both types of experiment, we grew the elutriated cells in 300 ml in a 1 l flask in a shaking incubator (300 rpm, 30 °C). At each sampling time point, we collected one sample to determine the population cell volume distribution with a CASY instrument. We collected a second sample, which we fixed in paraformaldehyde solution and kept for microscopy analysis to determine cell cycle phase distributions.

During the proteomics experiment we also collected a third sample for mass spectrometry proteomics. We collected a sample in a 50 ml tube and added ice-cold TCA to a final concentration of 10%. We then pelleted the cells by centrifuging (10 min, 3900 rpm, 4 °C) (Eppendorf, 5810 R), poured off the supernatant, and washed the pellet once with ice-cold PBS. We then flash-froze the pellet in liquid nitrogen and stored it at -70 °C until further processing for proteomics analysis.

### Cell volume distribution measurements

We took 50-100 µl of the synchronized culture and mixed it with 10 ml of CASYton (Omni Life Science, #5651808) in a CASYcup (Omni Life Science, #5651794) by inverting 10 times. We then performed measurements with a CASY TT (Omni Life Science) equipped with the 60 µm capillary, using a 400 µl sample volume. We exported data to a Microsoft Excel spreadsheet using CASYworkX software (Omni Life Sciences, version 1.26) to obtain the count values for each of the 400 particle diameter bins present in the CASY output. Each bin has a width of 0.05 µm and we took the middle value of a bin’s diameter range as the representative diameter for the bin *d*_*bin*_, e.g. for a bin with edges at 0.05 and 0.1 µm, the representative value for the bin was 0.075 µm. We estimated representative volumes for each bin, *v*_*bin*_ assuming spherical particles:

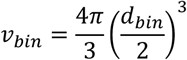

We performed additional filtering, baseline correction and normalisation using custom Python code implemented in the CASY class [see “Code availability”]. We removed counts originating from particles outside the normal size range for *S. cerevisise* by excluding any bins corresponding to particle diameters smaller than 2 µm or larger than 6 µm. We then corrected data from each experiment separately to remove a consistent baseline background count distribution which we observed in our datasets. To do so, for each time point we computed the fraction of the total counts present in each bin. Then, for each bin, we found the mean fraction of counts present in that bin across all time points in the experiment. We estimated a fractional count background across bins using an asymmetric least squares algorithm, using smoothing parameter lambda = 20000, as implemented in the pybaselines library (Erb; Eilers, 2003; Eilers & Boelens, 2005) (version 1.1.0). For each time point, we estimated the number of counts arising from the background by multiplying the estimated fractional count background for each bin by the total counts for the respective time point. We then subtracted those estimated background counts from the measured values and re-computed the fraction of total counts present in each bin at each time point, thus providing the normalised cell volume distribution.

### Fluorescence microscopy and analysis of formaldehyde-fixed cells

In the elutriation experiments used to test the yeast population model, to fix cells for imaging, we pelleted cells in 1 ml of culture using a benchtop centrifuge (accuSpin Micro 17R, Fisherbrand) (13000 rpm, 3 min, 4 °C), discarded the supernatant, resuspended the pellet in 100 µl of ice-cold formaldehyde solution (4% paraformaldehyde, 3.4% sucrose) and kept the sample on ice for 5 minutes. Then, we added 1 ml of ice-cold PBS and stored the sample at 4 °C until imaging. In the elutriation experiments used for proteomics, the method differed in that cells were initially pelleted for 30s prior to addition of formaldehyde solution.

Before imaging, we transferred 2–4 µl of formaldehyde-fixed cells onto a cover slip and placed a small rectangular of agar pad on top. For the experiments used to test the yeast population model, the pad was 2% agarose in YNB, but 2.3% agarose in PBS for the proteomics experiments. We imaged cells using a Nikon Eclipse Ti-E widefield inverted fluorescence microscope fitted with using an Andor DU-897EX camera. We used a 100x Nikon CFI Plan Apo λD Oil (NA = 1.45) objective for the experiments used to test the population model, and a 100x Nikon CFI Super Fluor Oil (NA = 0.5–1.3) for the proteomics experiments. We used a Lumencor AURA excitation source. We operated the microscope using Nikon NIS-Elements software.

For brightfield imaging, we illuminated cells with a halogen lamp, fitted with a 420 nm beam splitter to filter out UV wavelengths. For the experiments used to test the yeast population model, we used a 300 ms exposure, but a 100 ms exposure for the proteomics experiments.

To image sfGFP, we excited samples at 470 nm using a 470/40 nm band-pass filter, a 495 nm beam splitter, and a 525/50 nm excitation filter. For the experiments used to test the yeast population model, we used a light source intensity of 12% over a 50 ms exposure. For the proteomics experiments we used an intensity of 3% over a 200 ms exposure.

To image mCherry, we excited samples at 565 nm using a 560/40 nm band-pass filter, a 585 nm beam splitter, and a 630/75 nm emission filter. For the experiments used to test the yeast population model, we used a light source intensity of 25% over a 300 ms exposure. For the proteomics experiments, we used an intensity of 10% over 250 ms, 400 ms, and 900 ms for experiments 1, 2, and 3 respectively.

Per time point, we took at least 100 images and recorded data for at least 150 cells. We classified cells into one of five cell cycle phases (early-G1, late-G1, S/G2/M, anaphase or telophase) based on brightfield, Whi5-mCherry and histone Hta2-sfGFP appearance [Fig. 2B]. We used the following criteria to determine a cell’s cycle phase: cells in early-G1 were unbudded and showed a Whi5-mCherry focus coincident with a single nuclear Hta2-sfGFP focus. Cells in late-G1 were unbudded, had no Whi5-mCherry focus and had a single Hta2-sfGFP focus. Cells in S/G2 were budded, had no Whi5-mCherry focus and had a single Hta2-sfGFP focus. Cells in anaphase were budded, had no Whi5-mCherry focus and had either two Hta2-sfGFP foci or a single elongated or two-lobed one (indicative of karyokinesis having started). Finally, cells in telophase were budded, had two Whi5-mCherry foci and two Hta2-sfGFP foci. After counting the number of cells in each cell cycle phase, we divided by the total number of cells across all phases to obtain the fraction of the population per cycle phase.

### Proteomics sample preparation

We lysed 2×10^8^ cells in in 50 µl lysis buffer (2 M Guanidinium-HCl (Gua), 0.1 M ammonium bicarbonate, phosphatase inhibitors (Sigma P5726 & P0044)) by sonication (Bioruptor, 10 cycles, 30 seconds on/off, Diagenode, Belgium) and digested proteins as described previously (Ahrné *et al*, 2016). Briefly, we reduced proteins with 5 mM TCEP for 10 min at 95 °C and alkylated them with 10 mM chloroacetamide for 30 min at 37 °C. After diluting samples with 100 mM ammonium bicarbonate buffer to a final Gua concentration of 0.5 M, we digested proteins by incubation with sequencing-grade modified trypsin (1/50, w/w; Promega, Madison, Wisconsin) for 12 h at 37°C. After acidification using 5% TFA, we desalted peptides using C18 reverse-phase spin columns (Macrospin, Harvard Apparatus) according to the manufacturer’s instructions, dried them under vacuum and stored them at -20°C until further use.

We labelled sample aliquots comprising 12.5 μg of peptides with isobaric tandem mass tags (TMTpro 16-plex, Thermo Fisher Scientific). We resuspended peptides in 10 μl labelling buffer (2 M urea, 0.2 M HEPES, pH 8.3) by sonication, added 2.5 μl of each TMT reagent to the individual peptide samples, and incubated them for 1 h at 25°C shaking at 500 rpm. To control for ratio distortion during quantification, we added a peptide calibration mixture consisting of six digested standard proteins mixed in different amounts to each sample before TMT labelling (for details see (Ahrné *et al*, 2016)). To quench the labelling reaction, we added 0.75 μl aqueous 1.5 M hydroxylamine solution and incubated samples for another 5 min at 25°C with shaking at 500 rpm. We then pooled the samples, increased the pH of the sample pool to 11.9 by adding 1 M phosphate buffer (pH 12), and incubated for 20 min at 25°C with shaking at 500 rpm to remove TMT labels linked to peptide hydroxyl groups. After the incubation, we stopped the reaction by adding 2 M hydrochloric acid until a pH < 2 was reached. Finally, we further acidified peptide samples using 5 % TFA, desalted them using Sep-Pak Vac 1cc (50 mg) C18 cartridges (Waters) according to the manufacturer’s instructions, and dried them under vacuum.

We fractionated TMT-labeled peptides by high-pH reversed phase separation using a XBridge Peptide BEH C18 column (3,5 µm, 130 Å, 1 mm x 150 mm, Waters) on an Agilent 1260 Infinity HPLC system. We loaded peptides onto the column in buffer A (20 mM ammonium formate in water, pH 10) and eluted using a two-step linear gradient from 2% to 10% in 5 minutes and then to 50% buffer B (20 mM ammonium formate in 90% acetonitrile, pH 10) over 55 minutes at a flow rate of 42 µl/min. We monitored elution of peptides with a UV detector (215 nm, 254 nm) and collected a total of 36 fractions. We pooled these into 6 fractions using a post-concatenation strategy as previously described (Wang *et al*, 2011) and dried them under vacuum.

### Proteomics LC-MS analysis

We resuspended dried peptides in 0.1% aqueous formic acid and subjected them to LC–MS/MS analysis using a Q Exactive plus Mass Spectrometer fitted with an EASY-nLC 1000 (both Thermo Fisher Scientific) and a custom-made column heater set to 60°C. Peptides were resolved using a RP-HPLC column (75μm × 30cm) packed in-house with C18 resin (ReproSil-Pur C18–AQ, 1.9 μm resin; Dr. Maisch GmbH) at a flow rate of 0.2 μl min^-1^. We used the following gradient for peptide separation: from 5% B to 15% B over 10 min to 30% B over 60 min to 45 % B over 20 min to 95% B over 2 min followed by 18 min at 95% B. Buffer A was 0.1% formic acid in water and buffer B was 80% acetonitrile, 0.1% formic acid in water.

We operated mass spectrometer in DDA mode with a total cycle time of approximately 3 s. Each MS1 scan was followed by high-collision-dissociation (HCD) of the 10 most abundant precursor ions with dynamic exclusion set to 30 seconds. For MS1, 3e6 ions were accumulated in the Orbitrap over a maximum time of 100 ms and scanned at a resolution of 70,000 FWHM (at 200 m/z). MS2 scans were acquired at a target setting of 10^5^ ions, maximum accumulation time of 110 ms and a resolution of 35,000 FWHM (at 200 m/z). Singly charged ions and ions with unassigned charge state were excluded from triggering MS2 events. We set the normalized collision energy to 30%, the mass isolation window to 1.1 m/z, and one microscan was acquired for each spectrum.

### Mass spectrometry data analysis

We analysed the raw acquisition files using the SpectroMine software (Biognosis AG, Schlieren, Switzerland). Spectra were searched against a yeast database consisting of yeast protein sequences (downloaded from Uniprot on 2020/04/21) and 392 commonly observed contaminants (in total 12940 entries). We used Standard Pulsar search settings for TMT 16 pro (“TMTpro_Quantification”) exported the resulting identifications and corresponding quantitative values on the PSM level using the “Export Report” function. We used the acquired reporter ion intensities in the experiments for automated quantification and statistical analysis using our in-house developed SafeQuant R script (v2.3, (Ahrné *et al*, 2016)). This analysis included adjustment of reporter ion intensities, global data normalization by equalizing the total reporter ion intensity across all channels and summation of reporter ion intensities per protein and channel.

### Modelling yeast population growth

The computational model to simulate the temporal development of a partially synchronised yeast population in liquid culture in terms of cell volume and cell cycle phase was setup as follows. A simulated population consisted of individual simulated cell cycles, each of which was defined by a set of unique parameter values including cell cycle durations, an initial volume, and linear volume growth rates. As such, one cell cycle represented the volume growth and cell cycle progression of a single cell between two cytokinesis events. We distinguished between daughter cycles, i.e. the first cell cycle undergone by a cell after separating from its mother, and mother cycles, i.e. any subsequent cell cycle. The process of running a simulation corresponding to a given elutriated experimental population proceeded in two stages. First, we generated the initial set of cell cycles, i.e. the first cell cycles of the daughter cells obtained by elutriation. Second, we simulated the process of population growth by cell division, occurring over the duration of an experiment, by adding a subsequent mother and daughter cycle whenever a cycle in the population ended (i.e. reached cytokinesis) prior to the maximum simulation time.

Each cell cycle in a simulated population was fully described by the following 13 parameter values and a simulated population consisted of a matrix where each row contained parameter values for one cell cycle (for cell cycle event definitions, see section “Calculation of cell cycle parameter distributions” and Fig. 2B; for an illustration of a subset of these cycle parameters, see Fig. 3A).

1. Cycle ID: a unique integer value.
2. Previous cycle ID: the integer cycle ID of the immediately preceding cell cycle (for cycles generated by a cell division during simulation).
3. Generation number: an integer value recording how many division cycles that the cell, which a given cell cycle corresponded to, had undergone. This was 0 for daughter cycles and ≥ 1 for mother cycles.
4. Initial volume (fl), *v*_0_: the volume that the cell, which a given cycle corresponded to, had at the beginning of the cycle, i.e. when the previous cycle reached cytokinesis.
5. Initial time (min), *t*_0_: the point on the simulation timeline at which a cell cycle began, i.e. the time at which the previous cycle reached cytokinesis.
6. Early-G1 duration (min), *d*_1_: the duration between the cycle’s initial time and the time at when it reached START.
7. Late-G1 duration (min), *d*_2_: the duration between when a cycle reached START and when it reached budding.
8. S/G2 duration (min), *d*_3_: the duration between when a cycle reached budding and when it reached karyokinesis.
9. Anaphase duration (min), *d*_4_: the duration between when a cycle reached karyokinesis and when it reached mitotic exit.
10. Telophase duration (min), *d*_5_: the duration between when a cycle reached mitotic exit and when it reached cytokinesis.
11. Separation duration (min), *d*_6_: the duration between when a cycle reached cytokinesis and when the subsequent mother and daughter cells were no longer connected.
12. G1 linear volume growth rate (fl min^-1^), *g*_1_: the linear volume growth rate between birth/cytokinesis at the beginning of a cycle and budding.
13. S/G2/M linear volume growth rate (fl min^-1^), *g*_2_: the linear volume growth rate between budding and cytokinesis.

We simulated the development of a population from a simulation start time, *T*_0_, which corresponded to the completion of elutriation, up to a maximum time, *T*_*max*_, which was usually the final sampling time point in a time course. We considered a cycle to be ‘active’ at any simulation time, *T*, where *t*_0_ ≤ *T* ≤ *t*_*cyt*_. For a given active cycle, we could determine the cell volume, *v*,as:

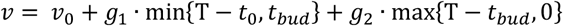

where *t*_*bud*_ = *t*_0_ + *d*_1_ + *d*_2_, and *t*_*cyt*_ = *t*_0_ + *d*_1_ + *d*_2_ + *d*_3_ + *d*_4_ + *d*_5_. As such, the minimum volume for a cycle was *v*_0_ at *t*_0_, the volume was *v*_*bud*_ when it reached budding at *t*_*bud*_, and the maximum volume was *v*_*cyt*_, achieved when it reached cytokinesis at *t*_*cyt*_. Similarly, we could determine the cell cycle phase of an active cycle at *T* by finding the latest incomplete cell cycle phase. For example, for a cell in early-G1, the inequality *t*_0_ ≤ *T* < *t*_0_ + *d*_1_ would hold. Similarly, if a cell was in late-G1, *t*_0_ + *d*_1_ ≤ *T* < *t*_0_ + *d*_1_ + *d*_2_ would hold instead, and so on for the other cell cycle phases.

When simulating a population, we first generated cell cycles corresponding to the first (daughter) cycles of the cells obtained by elutriation, numbering *n*_*initial*_. As part of this process, we ensured that the cell volume distribution of the simulated population matched the one measured for the corresponding experimental population. We measured the experimental cell volume distributions using a CASY instrument, whose output consisted of a set of volume bins and counts for the number of measured cells which fell into each volume bin. After processing the CASY measurements and filtering to exclude excessively large and small volume bins (see “Cell volume distribution measurements”), we obtained a count fraction, *f*_*bin*_, for each of the included volume bins. We generated parameter values (*v*_0_, *d*_1_, *d*_2_, *d*_3_, *d*_4_, *d*_5_, *d*_6_, *g*_1_, *g*_2_) for a pool of potential daughter cell cycles (numbering *n*_*pool*_) by sampling randomly from multivariate log-normal distribution D, computed from data obtained in microfluidics imaging experiments (see “Calculation of cell cycle parameter distributions”]. Then, for each volume bin in the CASY data, we determined the set of potential cycles (if any) satisfying *v*_0_ ≤ *v*_*bin*_ ≤ *v*_*cyt*_. From that set of cycles, we sampled (with replacement) a number of cycles corresponding to the fraction of *n*_*initial*_ equal to the CASY count fraction for the volume bin, i.e. *n*_*initial*_ **·** *f*_*bin*_. Then, we set *t*_0_ for each of those cycles such that, at simulation *T*_0_, they reached the volume at the midpoint of the volume bin edges. Having performed this for each volume bin, we assigned unique cycle IDs to all cycles and set their generation number to 0. In this way, we constructed an initial population of daughter cell cycles such that the volume distribution at *T*_0_ closely matched the CASY measurements from the experimental population bin-for-bin [Fig. 3B].

Next, we continued the development of the simulated population until *T*_*max*_ with a simulated cell division process whenever a cell cycle reached cytokinesis. At each cell division event, we created a new daughter cycle and a new mother cycle. For both new cycles, we generated unique cycle IDs, set their previous cycle ID to the cycle ID of the preceding cycle, and set *t*_0_ to *t*_*cyt*_ from the preceding cycle. For the daughter cycle, we set the generation number to 0, set *v*_0_ = *v*_*cyt*_ − *v*_*bud*_ from the preceding cycle, and sampled the remaining parameters (*d*_1_, *d*_2_, *d*_3_, *d*_4_, *d*_5_, *d*_6_, *g*_1_, *g*_2_) from joint log-normal distribution D conditioned on *v*_0_. For the mother cycle, we incremented the generation number from the preceding cycle, set *v*_0_ = *v*_*bud*_ from the preceding cycle, and sampled the remaining parameters (*d*_1_, *d*_2_, *d*_3_, *d*_4_, *d*_5_, *d*_6_, *g*_1_, *g*_2_) from joint log-normal distribution M conditioned on *v*_0_ [Fig. 3C]. As such, generation of a simulated population was complete once every cycle either finished after *T*_*max*_ or was followed by two subsequent cycles.

The Python functions for generating a simulated population are part of the Model class in the model.py source code file [see Code availability].

### Fitting cell population model log-normal distribution parameters

For each simulated population, we maximised the similarity to the corresponding experimental population by fitting the location and scale parameters of the joint log-normal distributions D and M. To perform fitting, we used a bounded iterative particle swarm optimisation algorithm (Kennedy & Eberhart) with Python code adapted from https://github.com/tisimst/pyswarm. Each particle in a swarm was defined by a set of 20 values, the 10 location and 10 scale parameters defining the constituent marginal log-normal distribution of the joint distributions D and M. At the start of an optimisation run, we generated values for each particle randomly by sampling uniformly between the bounds we selected for each scale or location parameter (see below). We also randomly selected a “velocity” vector for each particle through parameter space. At each iteration, for each particle we simulated a population using the particle’s distribution parameter values. We then evaluated the fit between the simulated population’s volume distributions and cell cycle phase distributions and the corresponding values measured during the relevant time course experiment (see below). At the end of each iteration, we updated each particle’s velocity based partly on the global best particle position and partly on the best of each particle’s own past positions. Using those “velocities”, we updated the parameter values defining each particle and returned to the simulation step.

To determine the fit between a pair of simulated and experimental populations, it was necessary to extract measures of population volume distribution and cell cycle phase distribution from the simulated population that were comparable to the ones measured experimentally at the same time points. For the volume distribution, at each time point we calculated the volume of every cell present in the simulated population. Importantly, we had to account for the fact that mother and daughter cells do not immediately separate following cytokinesis and so our CASY cell counter recorded such pairs as a single higher-volume particle rather than two discreet ones. As such, for time point *T*, whenever we calculated the volume of a mother and daughter pair, we combined the values into a single larger volume if 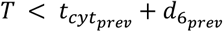, where 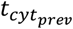 and 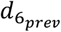 respectively are the cytokinesis time and separation duration from the preceding cell cycle. We then binned the simulated cell volumes, using the same bins as were used in the CASY output (see “Cell volume distribution measurements” section), and calculated the relative fraction of cell counts in each bin. We calculated the cell volume distribution component of the overall fit score as the sum of squared differences between the simulated and experimental relative bin counts at each time point.

To calculate the cell cycle phase distribution at each time point, we determined which phase (early-G1, late-G1, S/G2, anaphase or telophase) each simulated cell was in and then calculated the fraction of the total population in each phase. We calculated the cell cycle phase distribution component of the overall fit score as the sum of squared differences between the simulated and experimental fractions in each phase at each time point. The total fit score was the sum of the cell volume distribution, cell cycle phase distribution, and relative cell count components.

When performing fitting for the simulated population corresponding to a given experiment, we used a swarm size of 100 particles and allowed the swarm to converge over 25 iterations. We performed 100 repeats of the optimisation process with different starting values and selected the set of log-normal distribution parameter values which minimised the difference between the specific simulated and experimental populations. When generating simulated populations as part of the fitting process we used *n*_*initial*_ = 10^4^ and *n*_*pool*_ = 10^5^. When fitting simulated populations to experiments, we used the parameter bounds in Supp. Table 3.

The Python functions for fitting a simulated population to experimental data are part of the Model class in the model.py source code file [see Code availability].

### Definition of a standard cell cycle coordinate, *φ*

We defined a standard cell cycle progression coordinate, *φ*, a unitless coordinate with the domain [−1, 1], onto which any yeast cell at any point in its cell cycle could be mapped. Three cell cycle events correspond to fixed *φ* positions: a cell had coordinate *φ* = −1 at birth, i.e. the point of cytokinesis where it became a separate cytoplasmic compartment from its mother cell; a cell had coordinate *φ* = 0 at START; and cell had a coordinate of *φ* = 1 at the START in the subsequent cell cycle [Fig. 1A]. Thus, the region −1 ≤ *φ* < 0 on the *φ* axis represents the early-G1 phase of a daughter cell, which each cell progressed through only once. The regions 0 ≤ *φ* < *φ*_*bud*_, *φ*_*bud*_ ≤ *φ* < *φ*_*kar*_, *φ*_*kar*_ ≤ *φ* < *φ*_*exit*_, and *φ*_*exit*_ ≤ *φ* < *φ*_*cyt*_ represent the cell cycle phases late-G1, S/G2, anaphase, and telophase respectively, which are the same for mother and daughter cells. Lastly, the region *φ*_*cyt*_ ≤ *φ* < 1 represents early-G1 for a mother cell. We computed *φ*_*bud*_, *φ*_*kar*_, *φ*_*exit*_, and *φ*_*cyt*_ during the calculation of convolution matrices from a set of simulated populations (see “Modelling yeast population growth” section). The *φ* position of a cell increases as a function of time and cell cycle progression, with the exception that as soon as a mother cell reached *φ* = 1 it was reset to *φ* = 0.

We represented the mean cycle–dependent single-cell concentration of a biomolecule *c* as function of *φ, c*(*φ*). Since *φ* = −1 and *φ* = *φ*_*cyt*_ both correspond to cytokinesis events, we set *c*(−1) = *c*(*φ*_*cyt*_). Similarly, since *φ* = 0 and *φ* = 1 both correspond to START, we defined that *c*(0) = *c*(1). We discretised the *φ* axis into *b* equally spaced bins to represent a cell cycle–dependent biomolecular concentration dynamic as a vector **c** ∈ ℝ^*b*^ with ***c*** ≥ **0**, where the *j*-th element of **c**, *c*_*j*_, was the mean of *c*(*φ*) for *φ* values within the *j*-th bin [Fig. 1B].

### Derivation of the convolution equation

To estimate a cell cycle–dependent biomolecular concentration dynamic, **c**, from population-level concentration measurements by deconvolution, we derived an equation relating **c** to the measured concentration of the same biomolecule, ^*m*^**c**, in a sample of cells. The population-averaged concentration, ^*m*^**c**, of a biomolecule of interest in a sample is the total abundance of that biomolecule across all cells in the sample divided by the total cell volume in the sample. A sample at a given time point contains *p* cells with a total cell volume of *v*_*total*_. The *d*-th cell in the sample has a concentration of the biomolecule of interest, *q*_*i*_, and a cell volume, *v*_*i*_, and thus an abundance of the biomolecule *a*_*i*_ = *q*_*i*_*v*_*i*_. Therefore, the measured population-averaged concentration is:

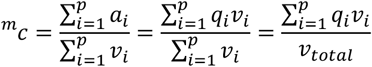

Each cell has a *φ* coordinate dependent on its cell cycle progression (see “Definition of a standard cell cycle coordinate, φ” section) and falls within one of the *b* bins on the discretised *φ* coordinate axis. From the definition of **c**, the mean single cell biomolecule concentration for cells in the *j*-th *φ* bin is the *j*-th element of **c**, *c*_*j*_. As such, we made the simplifying assumption that all cells within the *j*-th *φ* bin have biomolecule concentration *c*_*j*_. We also define *w*_*j*_ as the total volume of all cells within the *j*-th *φ* bin. Hence, we represented ^*m*^**c** as a summation over *φ* bins rather than individual cells:

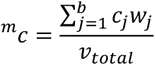

To simplify this equation, and express it in terms of linear algebra, we defined the vector **f** ∈ ℝ^*b*^ with **0** ≤ **f** ≤ **1**, where 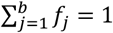, to be the fractions of total sampled cell volume comprised by cells within each *φ* bin. Here the *j*-th element of **f** is 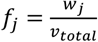 and hence we have:

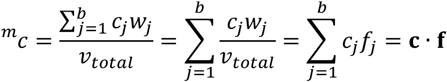

This equation represents a single population-level measurement. However, since vector **c** contains multiple elements, we required multiple measurements to solve for **c**. During a time course experiment, *n* measurements are made and hence the vector of measured population-averaged concentrations is ^*m*^**c** ∈ ℝ^*n*^ with ^*m*^**c** ≥ **0**. Let the matrix **F** ∈ ℝ^*n*×*b*^ with **0**_*n*×*b*_ ≤ **F** ≤ **1**_*n*×*b*_ where **F1**_*b*×1_ = **1**_*n*×1_ be the fractional distribution of sample cell volume across *φ* bins at each time point. We refer to **F** as the convolution matrix for a given experiment. With this definition, the expression representing the relationship between the population-level concentration measurements, ^*m*^**c**, and **c** is the matrix multiplication [Fig. 1D]:

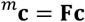

This equation models the ideal case where measured values are error free. We represent such errors by including an unknown vector error term **ε**:

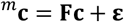

### Calculating a convolution matrix from a population model

To solve the convolution equation using measured concentration values from an experiment, we needed an accurate estimate of the convolution matrix **F** for that experimental population. To estimate a convolution matrix for an experiment, we used a simulated population that we generated using cell cycle parameter distribution parameters fitted to the corresponding experiment. When generating populations for subsequent calculation of convolution matrices, we used *n*_*initial*_ = 10^5^ and *n*_*pool*_ = 10^6^. To calculate the convolution matrix from a simulated population, for every sampling time point *T*, we determined the set of cells present in the population as those corresponding to the active cycles (i.e. cycles where *t*_0_ ≤ *T* ≤ *t*_*cyt*_). We then calculated the cell volume for each cell (see “Modelling yeast population growth” section); mapped each cell to a *φ* coordinate and hence to a *φ* bin; and then calculated the vector **f**, whose elements were the fraction of the total cell volume associated with cells in each *φ* bin. Finally, for an experiment with *n* time points, we stacked the *n* **f** vectors row wise to give the *n* × *b* convolution matrix **F**.

To map cells to *φ* coordinates based on their cell cycle phase, we first calculated values for *φ*_*bud*_, *φ*_*kar*_, *φ*_*exit*_, and *φ*_*cyt*_. We calculated the mean duration from budding until cytokinesis as the sum of the mean durations of late-G1, S/G2, anaphase, and telophase for the *p*_*b*_ daughter cycles (i.e. ones with generation number 0) as:

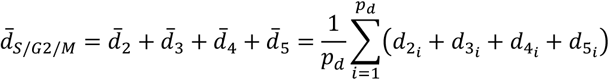

where 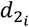 is the *d*_2_ cell cycle parameter for the *d*-th cell and so on. We also calculated the mean mother early-G1 duration from the *p*_*m*_ mother cycles (i.e. one with generation number > 0) as:

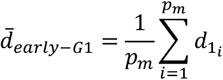

Hence, we calculated the average duration from START to subsequent START – equivalent to the *φ* region 0 ≤ *φ* < 1 as:

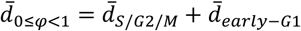

Finally, we calculated the *φ* coordinates *φ*_*bud*_, *φ*_*kar*_, *φ*_*exit*_, and *φ*_*cyt*_ as:

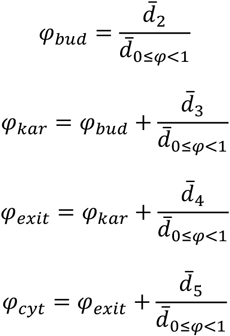

To perform deconvolution using data from multiple experiments together, we estimated convolution matrices using *φ*_*bud*_, *φ*_*kar*_, *φ*_*exit*_, and *φ*_*cyt*_ values averaged across the relevant experiments. To do so, we first determined *φ*_*bud*_, *φ*_*kar*_, *φ*_*exit*_, and *φ*_*cyt*_ from the simulated population from each experiment. Then we calculated the average *φ*_*bud*_, *φ*_*kar*_, *φ*_*exit*_, and *φ*_*cyt*_ values as the mean of the respective values across all replicate experiments. We used those mean values for subsequent calculations.

Having calculated *φ*_*bud*_, *φ*_*kar*_, *φ*_*exit*_, and *φ*_*cyt*_, we could map each cell to a *φ* coordinate at each time point. For a cell with an active cycle at a given time point *T*, we determined the current cell cycle phase and calculated, *ψ*, the fraction of that phase that the cell had completed. For a cell in early-G1, the fraction would be,

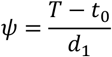

and for a cell in late-G1 the fraction would be,

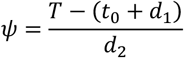

and so on for the other cell cycle phases.

We then mapped each cell to a corresponding *φ* coordinate using the positions of the key cell cycle events; we set a cell’s *φ* coordinate as the position of the cell cycle event preceding its current phase plus *ψ* times the distance to the following cell cycle event. For example, for a daughter cell in early-G1 the prior cell cycle event was birth/cytokinesis, defined as *φ* = −1, and the following event was START, defined as *φ* = 0, and so we calculated its *φ* coordinate as:

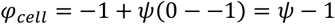

For a mother cell in early-G1, the prior event was cytokinesis, occurring at *φ*_*cyt*_, and the following event was the subsequent START, defined as *φ* = 1, and so we calculated its *φ* coordinate as:

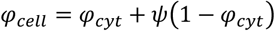

and so on for the other cell cycle phases.

Having mapped cells to *φ* coordinates, we assigned cells to the corresponding *φ* bin. We calculated the total volume of all cells at the given time point and, for each of the *b* bins, calculated the fraction of that total volume comprised by cells within the respective bin. As such, for each experiment, we obtained *n* vectors of fractional cell volumes (one per time point) and stacked them row wise to give the *n* × *b* convolution matrix **F**. When deconvolving our mass spectrometry proteomics results, we divided the *φ* coordinate axis into *b* = 20 bins and we had *n* = 15 sampled time points.

### Deconvolution

To estimate a cell cycle-dependent biomolecular concentration dynamic, **c**, from population-level concentration measurements ^*m*^**c**, we had to solve the convolution equation ^*m*^**c** = **Fc** + **ε** (see “Derivation of the convolution equation” section). As this equation is ill-conditioned, we used an optimisation method to find a solution **ĉ**, which approximates **c**, by minimising a regularised least squares cost function. We used regularisation to control the tendency toward extremely rough solutions and did so by penalising a solution’s cost function dependent upon the roughness of the solution. We quantified the roughness of solution vector **c** as the sum of squared second order central finite differences over *φ*:

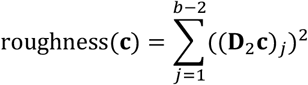

where the index *j* denotes the *j*-th bin of *φ*, (**D**_2_**c**)_*j*_ is the *j*-th element of the vector resulting from the matrix multiplication **D**_2_**c**, and **D**_**2**_ ∈ ℝ^*b*−2×*b*^ is the second order central finite difference matrix:

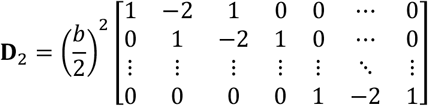

To control the influence of the roughness penalty on the overall cost function, we scaled the roughness penalty by *λ*, a regularisation coefficient. We also multiplied the roughness penalty by 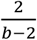 to normalise for both the *φ* bin width and number of second order central finite differences, such that the full roughness penalty for solution **c** is:

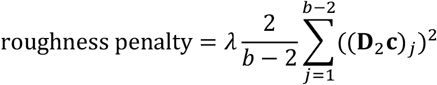

Hence, the complete, regularised least squares cost function to solve ^*m*^**c** = **Fc** + **ϵ** is,

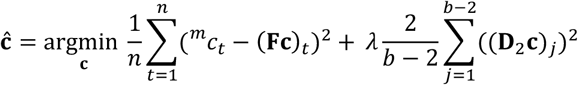

where the index *t* denotes the *t*-th measurement time point and (**Fc**)_*t*_ is the *t*-th element of the vector resulting from the matrix multiplication **Fc**. This equation has the analytic linear algebra solution [derivation in Supp. File 1]

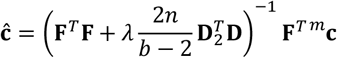

However, we were unable to use this analytic solution because the values we obtained from mass spectrometry proteomics were not direct measurements of concentration, meaning that ^*m*^**c** ≠ **Fc**. Instead, we had values which approximated relative concentration measurements, obtained by normalising mass spectrometry measurements for each protein within a single experiment by their mean (see “Mass spectrometry data analysis” section) that required that we further modify the cost function. Here we use the notation **z** to indicate the mean of the values of any vector **z** ∈ ℝ^*b*^ where:

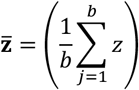

As such, we obtained relative measured concentration values ^*rm*^**c** for a single experiment as:

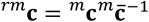

and sought to estimate the cell cycle–dependent relative biomolecular concentration dynamic ^*r*^**c** where:

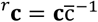

It was required that we solve ^*rm*^**c** = **F** ^*r*^**c** + ^*r*^**ε**, where ^*r*^**ε** denotes another unknown error term. We could find ^*r*^ĉ, an approximation of ^*r*^**c**, using the modified cost function:

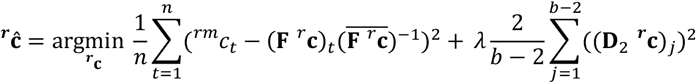

which we further adapted to facilitate the inclusion of data from *s* replicate time course experiments, providing the final multi-replicate cost function:

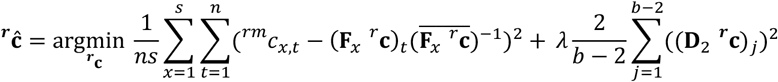

where ^*rm*^*c*_*x,t*_ is the relative measured concentration value at the *t*-th timepoint for the *x*-th replicate and **F**_*x*_ is the convolution matrix for the *x*-th replicate.

We estimated ^*r*^ĉ for each protein in our dataset separately, using data from our three replicate proteomics experiments and the corresponding convolution matrices estimated from fitted population models (see “Calculating a convolution matrix from a population model” section). During the solution process, we calculated an initial guess ^*r*^ĉ_*initial*_ using an adaptation of the analytic solution to ^*m*^**c** = **Fc** + **ε**:

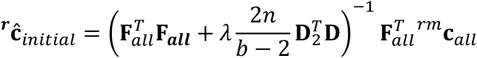

where **F**_***all***_ is the result of row wise concatenation of all three convolution matrices and ^*rm*^**c**_*all*_ is the result of row wise concatenation of all three sets of relative measured protein concentration values. We then used ^*r*^ĉ_*initial*_ as the starting value for ^*r*^**c** in a bounded optimisation of the multi-replicate cost function with the lower bound ^*r*^**c** ≥ (0.1)**1** and no upper bound. We also applied constraints such that solutions satisfied 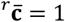 and that they were consistent with the original definition of the continuous function *c*(*φ*) (see “Definition of a standard cell cycle coordinate, φ” section). Specifically, we required that *c*(−1) = *c*(*φ*_*cyt*_) and *c*(0) = *c*(1). Since ^*r*^**c** is discreet, with values corresponding to the mean values within *φ* bins, we estimated the values at specific *φ* coordinates either by linear interpolation between bin midpoints or linear extrapolation of the terminal bin values.

To determine an appropriate value of the regularisation coefficient *λ* separately for each protein, we used a leave-one-out cross validation method [Fig. 6C]. Firstly, we defined the range of possible *λ* values as {10^−1^, 5 × 10^−2^, 10^−3^, 5 × 10^−3^, 10^−4^, 5 × 10^−4^, 10^−5^, 5 × 10^−5^, 10^−6^, 5 × 10^−6^, 10^−7^}. For each of these *λ* values, we excluded the measured data from one replicate and solved for ^*r*^**c** using only the data from the remaining two replicates. We then convolved that solution with the convolution matrix from the excluded replicate, divided that convolved solution by its mean, and found the residuals between that mean-normalised convolved solution and the relative concentration measurements from that excluded replicate. We repeated this process, leaving out each replicate in turn, and then calculated the mean squared error from all the residuals for the current λ value. We repeated this process for every possible *λ* value and selected the *λ* value which resulted in the minimal mean squared error [Fig. 6D]. We then performed a final deconvolution for the current protein, using the selected *λ* value and relative measured concentration data for all three replicates, to solve for ^*r*^**c**.

The Python functions for performing the deconvolution process are part of the Deconvolver class in the deconvolution.py source code file [see Code availability]. We used the “SLSQP” (sequential least squares programming) method of the minimize function from the scipy.optimize module (version 1.16.0) to perform the bounded optimisation with constraints.

### Filtering of deconvolved solutions

We performed deconvolution for 3373 proteins and filtered the deconvolved solutions to reduce the number of false positives, i.e. solutions containing spurious dynamics, and to enrich for the most dynamic solutions. To construct an appropriate filter for false positives, we assumed that deconvolution would be most accurate when the variation in measured relative concentration data for a protein had a higher ‘signal’ (i.e. variance due to cell cycle–dependent processes) than ‘noise’ (i.e. variance from error processes). As such, we estimated a signal-to-noise ratio metric and used it as a proxy for the plausibility of a deconvolved solution. For a given protein, we estimated the hypothetical noise-free mass spectrometry measurements by re-convolving the solution obtained from deconvolution. We then divided by the mean, centred these values around zero, and calculated the “signal” by summing the squared values, i.e.

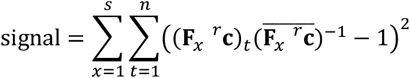

We estimated the ‘noise’ for a given protein as the sum of squared residuals between the normalised measured data and the normalised hypothetical noise-free mass spectrometry measurements, as

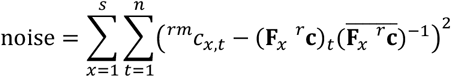

To additionally filter for proteins with the most dynamic solutions, we also calculated the peak-to-trough ratio of a deconvolved solution as:

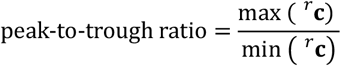

We selected the 563 proteins which were in the upper quartile for both “signal”-to-noise ratio and peak-to-trough ratio for downstream analysis. We refer to this as our “higher confidence subset”.

### Hierarchical clustering and GO enrichment analysis

We performed hierarchical clustering of the 547 filtered deconvolved solutions using the “complete” linkage method and taking the Pearson correlation between deconvolved solutions as the distance metric. We cut the cluster dendrogram such that proteins were classified into 10 different clusters. We chose this number as, with 10 clusters, deconvolved solutions with single-peak behaviours were clearly differentiated from solutions with higher frequency dynamics under visual inspection.

We used GO enrichment analysis, based on “Biological Process” annotation, to identify functional enrichment among the genes coding for the proteins assigned to each cluster, using the GOATOOLS Python library (Klopfenstein *et al*, 2018) (version 1.4.12). We obtained the general GO DAG file from https://current.geneontology.org/ontology/go-basic.obo on 2025/11/14 and the set of *S. cerevisiae* gene GO annotations from http://sgd-archive.yeastgenome.org/curation/literature/gene_association.sgd.gaf.gz on 2025/11/14. For all analyses, we included annotations with all evidence codes except for: No biological Data available (ND), Non-traceable Author Statement (NAS), and Inferred from Electronic Annotation (IEA). We used annotation from proteome accession “UP000002311” on https://www.uniprot.org (downloaded on 2025/08/18) to map between Uniprot protein accessions and *S. cerevisisae* genes. To test for statistically significant enrichment of GO terms within a given cluster, we used a two-sided Fisher’s exact test and corrected p-values for multiple testing with the Benjamini-Hochberg method, as implemented in the run_study method of the GOATOOLS GOEnrichmentStudy class. We used the genes corresponding to the 3373 proteins identified in our mass spectrometry dataset as the background gene set for enrichment testing.

We defined the “cytosolic ribosomal proteins” as the 105 proteins in our dataset with “Cellular Component” GO annotation GO:0022625 “cytosolic large ribosomal subunit” or GO:0022627 “cytosolic small ribosomal subunit”. We defined the “ribosome biogenesis proteins” as the 95 proteins in our datatset with “Biological Process” GO annotations GO:0042254 “ribosome biogenesis”, GO:0042273 “ribosomal large subunit biogenesis”, or GO:0042274 “ribosomal small subunit biogenesis”. We defined the “ergosterol biosynthesis enzymes” as the 21 proteins in our dataset which were annotated as part of the “superpathway of ergosterol biosynthesis I” on YeastPathways (https://pathway.yeastgenome.org/YEAST/NEW-IMAGE?type=PATHWAY&object=ERGOSTEROL-SYN-PWY-1).

### Transcription factor analysis

We obtained TF-gene interaction data with expression and DNA binding evidence from YEASTRACT (https://yeastract.com/) (Teixeira *et al*, 2023) on 2025/08/12. We downloaded TF-gene interaction matrices for 3370 genes whose protein products were in our proteomics dataset, treating activating and inhibiting regulatory relationships separately. In these two matrices, a TF gene interaction was represented by a 1 and no interaction by a 0. We merged the two TF-gene interaction matrices by subtracting the matrix representing inhibiting interactions from the matrix for activating ones, resulting in a matrix where a 1 indicated an activating relationship, a -1 indicated a repressive relationship, and 0 indicated either no relationship or ambiguous evidence. We filtered this TF-gene interaction matrix to only include TF-gene relationships for genes whose protein products were in our 563 higher confidence subset.

To quantify the amount of evidence consistent with an increase in activity of each TF across each successive pair of *φ* bins, we first calculated the log_2_ concentration fold change for each protein between each successive pair of *φ* bins. To avoid loss of resolution over the *φ* axis when calculating fold changes, we extended the deconvolved trajectories by one bin on the left and right side of the trajectory. Prior to cell birth at *φ* = −1, we inserted values from the bin with centre *φ* = 0.85, i.e. the bin immediately prior to cytokinesis. Following the second START at *φ* = 1, we inserted values from the bin with centre *φ* = 0.05, i.e. the bin immediately following the first START. Having calculated log_2_ fold changes, we normalised the values for each protein by dividing by the highest magnitude bin-to-bin log_2_ fold change undergone by that protein. We then determined the matrix product between the normalised log_2_ fold changes and the TF-gene interaction matrix. We normalised these TF activity-change scores by dividing the values for each TF by the number of its targets in the interaction matrix. Finally, we calculated TF activity scores for each TF by taking the cumulative sum of its TF activity-change scores across *φ* bins and then centered these scores by subtracting the mean for each TF separately. We summarised the magnitude of the overall dynamic for a given TF as the “activity score amplitude”, i.e. the minimum value subtracted from the maximum value.

To identify TFs with a higher-than-expected activity score amplitude, we used a Monte Carlo method. For each TF with *n*_*activated*_ activated targets and *n*_*inhibited*_ inhibited targets, we performed 10^5^ trials where we randomly sampled (without replacement) *n*_*activated*_ and *n*_*inhibited*_ proteins from our higher confidence subset. In each trial, we calculated the TF activity score amplitude as if those sampled proteins were the real activated and inhibited targets of the TF. We selected the 20 TFs for which the actual activity score amplitude was greater than at least 95% of the random trials for further analysis.

## Data availability

All mass spectrometry proteomics data associated with this manuscript have been deposited to the ProteomicsXchange consortium via MassIVE (https://massive.ucsd.edu) with the accession number MSV000098520 (Dataset is currently private. Reviewers can access the dataset following the instructions on the website. Username: MSV000098520_reviewer, Password: PCF). All other data is available on DataVerse https://doi.org/10.34894/AV0SDY.

## Code availability

The Python classes and functions used for population modelling and deconvolution are available on GitHub at https://github.com/molecular-systems-biology/Zylstra-et-al-2026.

The Jupyter notebooks containing Python code used to perform the analysis are available, along with the input and processed data, on DataVerse at https://doi.org/10.34894/AV0SDY.

For peer review purposes the data and code can be accessed via this temporary link to the DataVerse page: https://dataverse.nl/previewurl.xhtml?token=60346dee-87fa-429e-a15e-b99151830e56.

## Acknowledgements

Funding is acknowledged from the Dutch Research Council (NWO) (VICI, VI.C.192.003, to MH), (VIDI, 016.Vidi.189.116 to AMA) and from the European Union’s Horizon 2020 research and innovation programme (COFUND project oLife, grant agreement No 847675, to MH).

## Supplementary Figures

**Supplementary Figure 1.**
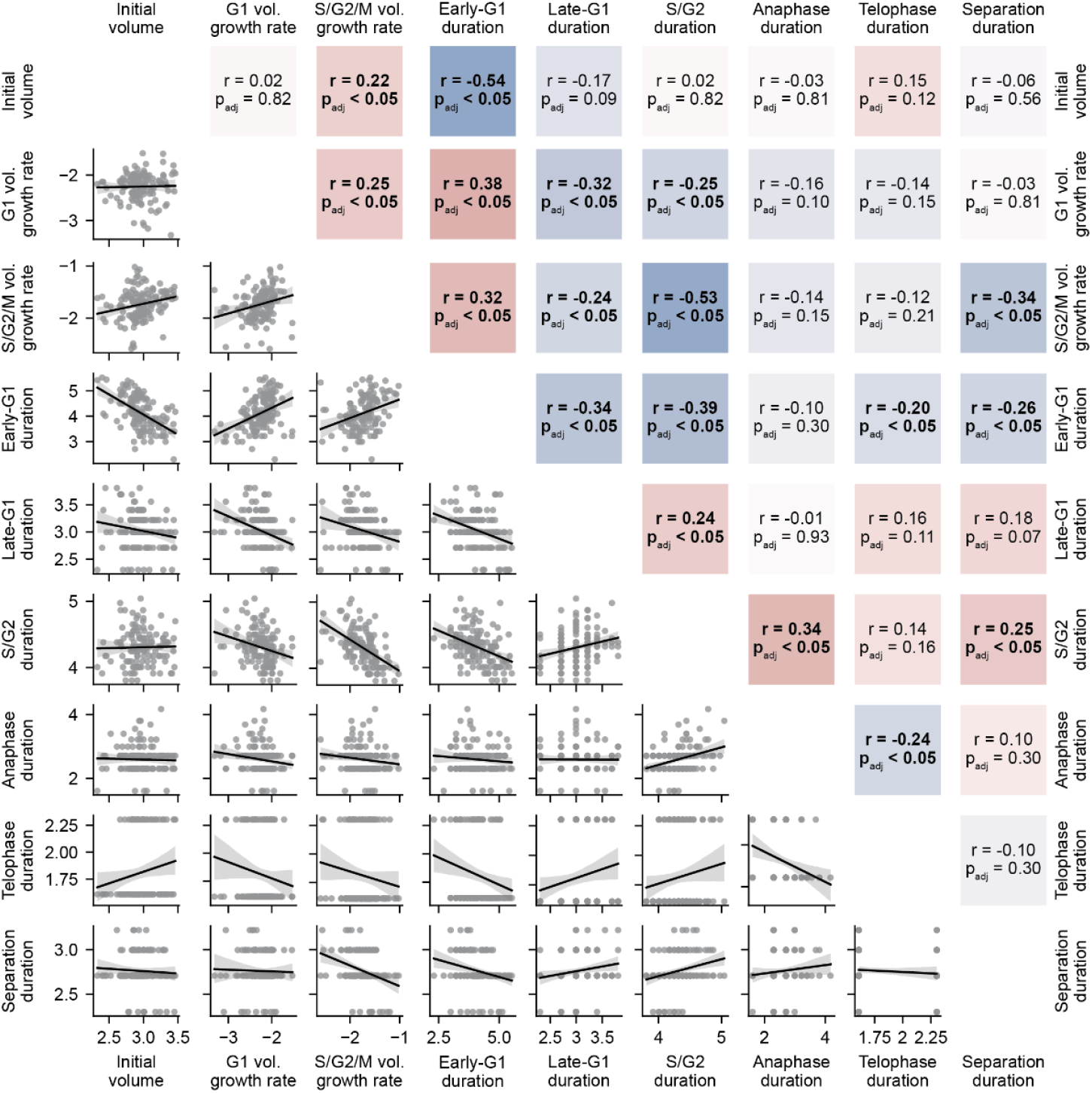
Comparison and Pearson correlations between log-transformed daughter cell cycle parameters. Lower triangle: black lines indicate linear regression; grey filled area indicates the 95% confidence interval of the mean. All plotted values are natural log–transformed. Upper triangle: r values indicate the pairwise Pearson product moment correlation coefficients between log-transformed parameters. Adjusted p-values were obtained using the Benjamini-Hochberg method to account for multiple testing. Squares with bold annotation text have p adj. < 0.05. Redder squares indicate positive correlations, bluer squares indicate negative correlations, colour intensity indicates correlation magnitude.

**Supplementary Figure 2.**
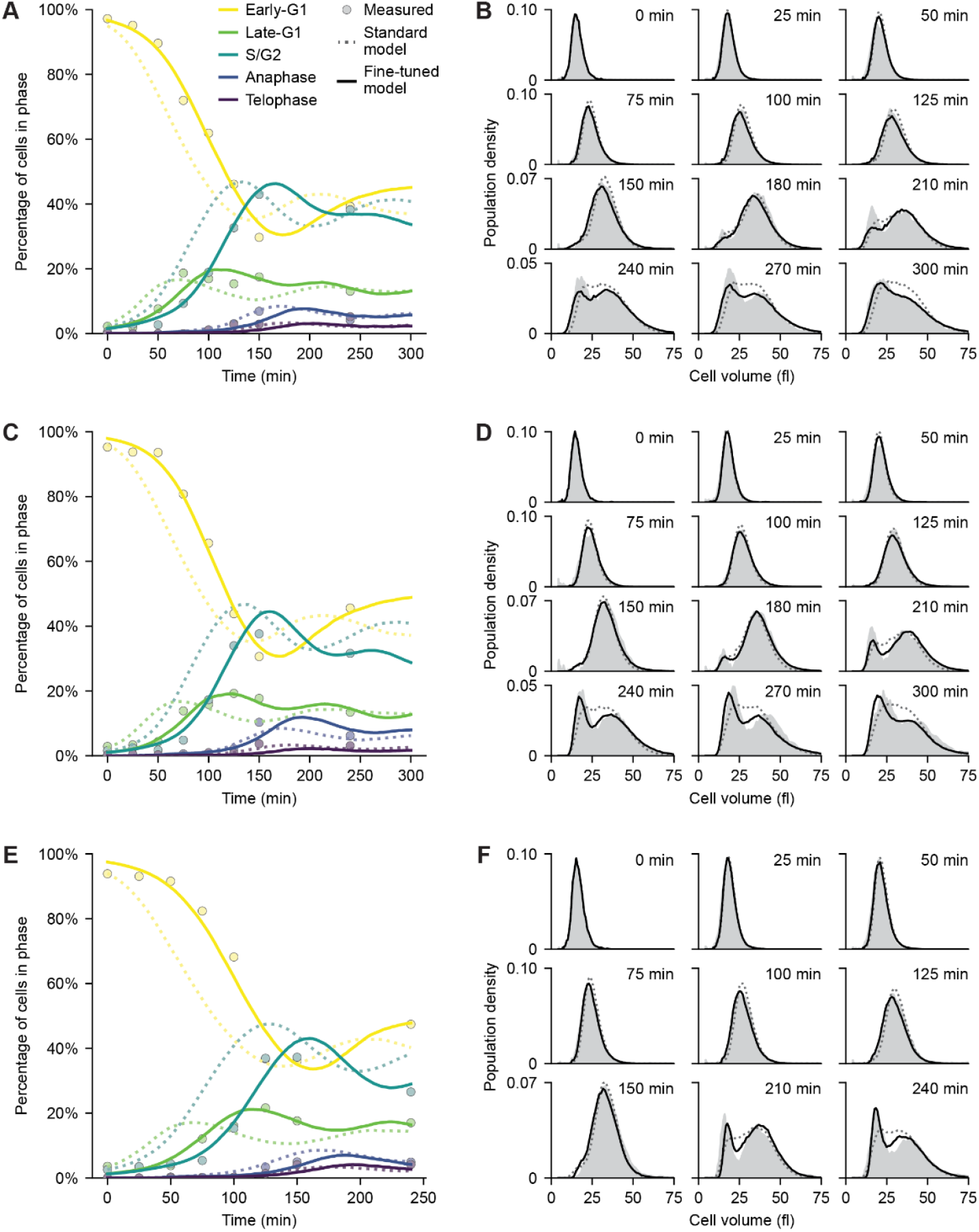
Full comparison between measured data, standard, and fine-tuned population model simulations for the three experiments used to test the population model predictions. **A**. Comparison of the measured cell cycle phase distributions of time course experiment 1with two corresponding simulated populations, generated either with parameter distributions as in Fig. 2 (Standard model) or parameter distributions fine-tuned to the specific experiment (Fine-tuned model). **B**. Comparison as in **A** for measured cell volume distributions at different time points (grey). Meaning of the lines, see legend in **A**. **C**. As in **A** for experiment 2. **D**. As in **B** for experiment 2. **E**. As in **A** for experiment 3. **F**. As in **B** for experiment 3.

**Supplementary Figure 3.**
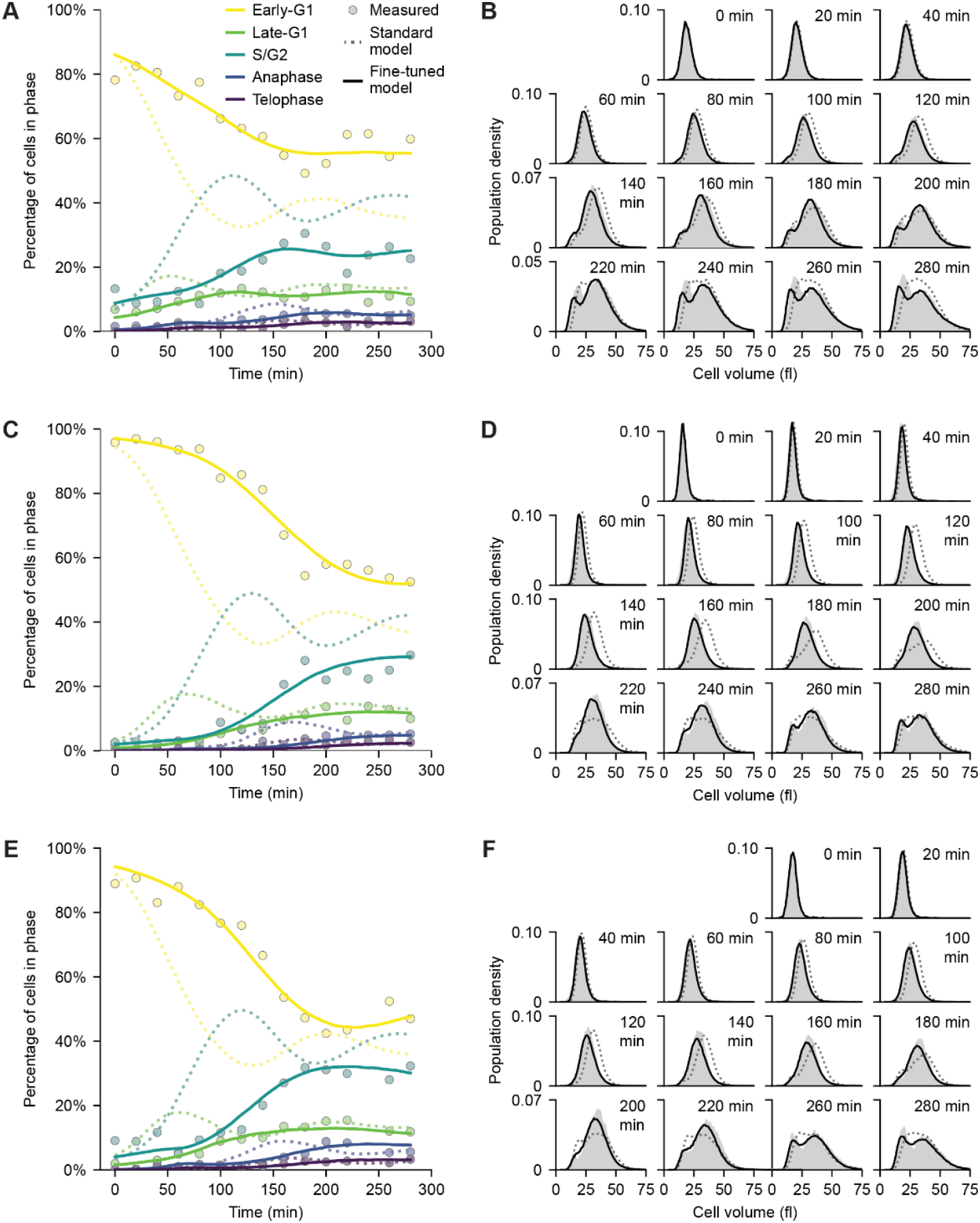
Full comparison between measured data, standard, and fine-tuned population model simulations for proteomics experiments. **A**. Comparison of the measured cell cycle phase distributions of proteomics time course experiment 1 with two corresponding simulated populations, generated either with parameter distributions as in Fig. 2 (Standard model) or parameter distributions fine-tuned to the specific experiment (Fine-tuned model). **B**. Comparison as in **A** for measured cell volume distributions at different time points (grey). Meaning of the lines, see legend in **A**. **C**. As in **A** for proteomics experiment 2. **D**. As in **B** for proteomics experiment 2. **E**. As in **A** for proteomics experiment 3. **F**. As in **B** for proteomics experiment 3.

**Supplementary Figure 4.**
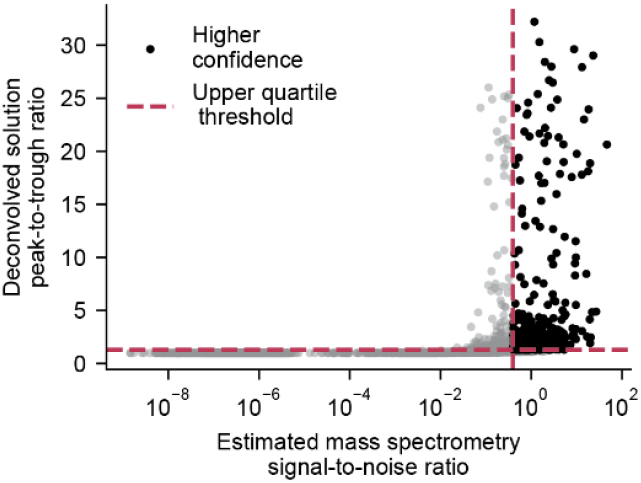
Selection of proteins with “higher confidence” deconvolved solutions. Two thresholds were used to filter for proteins with higher confidence deconvolved dynamics. For each protein, the hypothetical noise-free mass spectrometry measurements were estimated by re-convolving the solution obtained via deconvolution. The resultant values were divided by the mean, centred around zero and the “signal” calculated by summing the squared values. The noise was calculated as the summed squared residuals between the re-convolved solution and the measurement data. Deconvolved solution peak-to-trough ratios were calculated as the maximum value of a protein’s deconvolved solution divided by the minimum value. The subset of proteins with “higher confidence” deconvolved solutions was selected as the 563 proteins in the upper quartile for both estimated signal-to-noise ratio and peak-to-trough ratio.

